# Cell-specific dedifferentiation in *Arabidopsis thaliana*

**DOI:** 10.64898/2026.05.21.726904

**Authors:** Fabio Gomez-Cano, Luguang Jiang, Yoonjin Cho, Mariel Cruz-Gomez, Kook Hui Ryu, Kelsey M. Reed, Rachel Rivero, Lane Vitek, Bastiaan O.R. Bargmann, Alexandre P. Marand

## Abstract

Single somatic plant cells are capable of dedifferentiation and reversion to totipotency. However, understanding of dedifferentiation remains incomplete, limiting efforts to reprogram plant cell identities with engineered properties. Here, we describe the transcriptional dynamics of 125,091 *Arabidopsis thaliana* whole seedling protoplasts along a time course of dedifferentiation and varied chemical perturbations using single-cell RNA sequencing. We observed widespread cell-type-specific transcriptional reprogramming concomitant with cell wall digestion, including a guard cell-specific somatic embryogenic-like program initiated by ectopic *LEC2* expression. Contrasting cell states between chemical perturbations across real time revealed that phytohormones antagonize cell aging, promote retention of transcriptional diversity, and selectively suppress wound responses. By dissecting time-resolved transcriptional dynamics, we discovered that dedifferentiation is accompanied by the activation of stem-cell and meristematic marker genes and associated with prevalent chromatin remodeling. Despite discovery of a shared regulatory program underlying early dedifferentiation, temporal modeling of diverse somatic cell types uncovered extensive bias in dedifferentiation potential, including epithem-derived dedifferentiated cells, variable dedifferentiation rates, and diversity in early reprogramming outcomes as a major source of in vitro heterogeneity. Taken together, these data showcase how initial somatic cell identity, phytohormone signaling, chromatin remodeling, and reprogramming heterogeneity combine to coordinate the early transcriptional events defining cellular dedifferentiation.

## INTRODUCTION

In contrast to most animals, plant cells are uniquely plastic in their developmental potential; ∼1% of differentiated somatic plant cells retain the capacity to reprogram, re-enter the cell cycle, and regenerate a whole plant when cultured as protoplasts *in vitro*^1^. Regeneration is intrinsically related to wound response mechanisms – dissociation of protoplasts from their native tissue context is perceived autonomously as a major wounding event by cell wall integrity surveillance factors^2^. Past studies have revealed that wound and stress signals from cell wall integrity sensors rapidly activate calcium influx, reactive oxygen species production, and receptor-like and mitogen-activated protein kinase signaling, triggering large-scale transcriptional reorganization^3–5^. Several *APETALA2/ETHYLENE RESPONSE FACTOR* (*AP2/ERF*) transcription factors (TFs), such as *WOUND INDUCED DEDIFFERENTIATION 1* (*WIND1*), are among the earliest wound-induced TFs that act to promote dedifferentiation of somatic identity and establish partial competence for regeneration^6,7^. Dedifferentiation describes the repression of somatic identity transcriptional programs and is followed by the activation of factors associated with somatic embryogenesis or organogenic competency, such as *WUSCHEL* and *WUSCHEL-RELATED HOMEOBOX* (*WOX*) genes, *BABY BOOM* (*BBM*), *AGAMOUS-LIKE 15* (*AGL15*), and *ENHANCER OF SHOOT REGENERATION* (*ESR1/2*)^1,8–12^. Suppression of photosynthetic machinery, widespread chromatin remodeling, and ribosomal biogenesis are additional hallmarks of dedifferentiation in nascent protoplasts^13–17^. However, prior studies have focused on how dedifferentiation is controlled in mesophyll, limiting understanding of the impact of initial somatic cell identity.

In addition to transcriptional reprogramming, re-entry into the cell cycle is required for the formation of undifferentiated callus with organogenic potential^1,13,15,18,19^. Reactivation of cell proliferation is largely accomplished through the activity of phytohormones auxin and cytokinin. Cell cycle reinitiation in mesophyll protoplasts requires the activity of histone acetyltransferases that permit expression of root meristem identity *PLETHORA* (*PLT*) transcription factors, which in turn, induce expression of *YUCCA1* (*YUC1*)^15^. Auxin biosynthesis by *YUC1* triggers expression of *MYB-DOMAIN PROTEIN 3R4* (*MYB3R4*), a regulator of G2/M, through the *AUXIN RESPONSE FACTOR* (*ARF*) *7/9* and *AUXIN/INDOLE-3-ACETIC ACID INDUCIBLE* (*Aux/IAA*) *3*/*18* auxin signaling pathway^15^. However, auxin is necessary but not sufficient for cell cycle re-entry, and requires the phytohormone, cytokinin, to induce cell proliferation^20^. Similar to auxin, recent studies have demonstrated that cytokinin promotes cell cycle entry through MYB3R4 activation^21^. While autonomous phytohormone biosynthesis in protoplasts is sufficient to promote cell cycle re-entry, supplementation of exogenous auxin and cytokinin phytohormones significantly increases the frequency of cell proliferation in vitro^13^. In explants, cellular proliferation gives rise to pluripotent callus that develops both root and shoot stem cell potential depending on the ratio of auxin and cytokinin^22,23^, with root meristem identity typically acquired first by expression of *WOX*, *PLT*, and *LATERAL ORGAN BOUNDARY DOMAIN* (*LBD*) gene families^23–25^. Somatic embryogenesis via ectopic expression of *LEAFY COTYLEDONS 2* (*LEC2*) and other embryogenic factors also stimulates phytohormone biosynthesis and cell proliferation^26,27^. Past studies have found that somatic identity is a key factor in hormone-based transcriptional reprogramming, for example, with auxin inducing cell-type-specific gene expression changes in the Arabidopsis root^28^. However, how phytohormones affect cell fate trajectories during the early stages of dedifferentiation in protoplasts with diverse cellular identities remains unclear.

To address these gaps, we profiled the transcriptomes of 125,091 protoplasts derived from whole *Arabidopsis thaliana* seedlings along a six-day time course of in vitro culturing with and without exogenous phytohormones. We observed widespread transcriptional reprogramming following cell wall digestion and significant bias in somatic cell type representation across time. Analysis of temporal effects revealed that phytohormones attenuate cell age, favor expression of stem cell marker genes, promote transcriptional and cell state diversity, enhance proliferation, and repress a generalized wounding response. Although only a small fraction of protoplasts will ultimately have regenerative competency, we observed widespread and heterogeneous activation of stem cell marker genes in most profiled cells, including the unexpected cell-type-specific activation of factors involved in somatic embryogenesis immediately following tissue dissociation. Re-investigation of chromatin accessibility enabled the identification of regulatory factors and multiscale footprints associated with chromatin remodeling at the onset of dedifferentiation. Finally, mapping of time-resolved dedifferentiation trajectories for diverse cell types uncovered bias in dedifferentiation potential, variation in somatic reprogramming rates, and a generalized regulatory program defining the early stages of dedifferentiation across distinct somatic cells. Together, these data provide insight into the regulatory dynamics underlying the early stages of dedifferentiation conditioned by diverse somatic cell identities.

## RESULTS

### Cell wall loss induces widespread transcriptional reprogramming

To better understand protoplast phenotypes in culture, we isolated and imaged 11-day old *Arabidopsis thaliana* (Arabidopsis) seedling protoplasts carrying a nuclear (*p35S::H2B-RFP*) and cell membrane (*pUBQ10::myr-YFP*) reporter in liquid protoplast induction media^13^ with and without the addition of callus-promoting ratios of auxin and cytokinin phytohormones (2,4-dichlorophenoxyacetic acid [2,4-D] and 6-Benzylaminopurine [6-BA]; **Fig. 1a**; **Fig. S1a**). Cell budding and nuclei doubling became evident on Day 3, followed by cell morphology changes and evidence of cell division by Day 6 (**Fig. 1b**). Automated quantification of individual protoplast nuclei counts over a six day time course indicated that hormone treatments were associated with increased frequency of nuclei doubling (**Fig. 1c**-**1d**). Overall, protoplast phenotypes observed in our study were consistent with prior observations and suggestive of dedifferentiation^13^.

**Figure 1:**
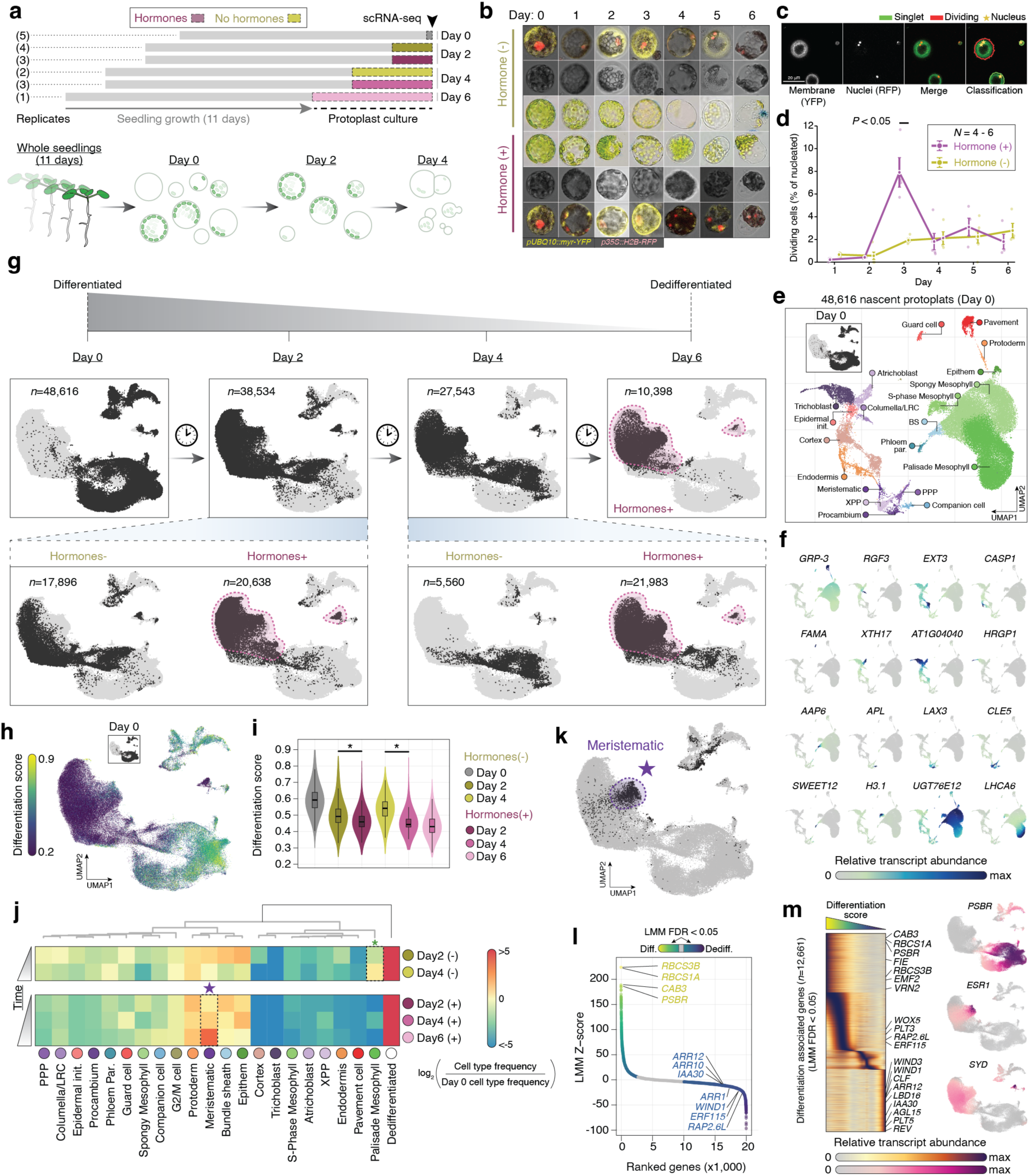
Widespread transcriptional reprogramming during dedifferentiation. (**a**) Experimental overview, design, sampling timeline, and number of biological replicates (**b**) Representative brightfield (center) and confocal (top and bottom) micrographs of nascent (Day 0, no treatments possible) and cultured protoplasts (Day 1-6) with and without auxin and cytokinin treatments. Red fluorescence indicates *p35S::H2B-RFP* expression and yellow indicates *pUBQ10::myr-YFP* expression. (**c**) Representative images from automated cell nuclei counting. (**d**) Time course of cell division frequency for hormone (purple) and non-hormone (yellow) treated cultures. (**e**) Uniform manifold approximation projection (UMAP) embeddings for Day 0 cells. Different colors reflect different cell type annotations. Inset indicates the same Day 0 cells in Fig. 1E. BS, bundle sheath; Init, initials; LRC, lateral root cap; Par, parenchyma; PPP, phloem pole pericycle; XPP, xylem pole pericycle. (**f**) Visualization of relative marker gene expression on the UMAP embedding (normalized expression values range 0-1 across cells for each marker gene). (**g**) UMAP embeddings for the fully integrated data set ordered by increasing dedifferentiation and sampling time. Day 0 cells (top left panel), Day 2 cells (top middle left panel), Day 4 cells (top middle right panel), and Day 6 cells (top right panel) are illustrated with black dots in each panel. UMAP embeddings for Day 2 and Day 4 cells split by hormone treatment are depicted in the bottom row (Day 2, left. Day 4, right). Cell populations (clusters 1, 2, 6, 7, 9, & 11) depleted of Day 0 cells and enriched in the phytohormone treatment from Day 2, 4, and 6 samples are showcased by a purple shaded outline. (**h**) Per-cell differentiation scores visualized on the UMAP embedding of the integrated data set. The red outline highlight cells with meristematic identity. (**i**) Distribution of differentiation scores among conditions. Asterix indicates Wilcoxon Rank Sum test *P-*value < 2.2e^-16^. (**j**) Heatmap illustrating the log_2_ fold-change in cell type frequencies between extended protoplast culturing (2+ days) and in planta (Day 0). The purple star and green asterisk highlight significant (FDR < 0.05) increases in meristematic and palisade mesophyll cell type proportions in phytohormone supplemented and deficient media, respectively. (**k**) UMAP embedding highlighting meristem-like cell identities. (**l**) Ranked distribution of Z-scores based on the linear mixed models (LMM) of differentiation score fit to normalized expression levels and technical covariates across cells for each gene. Non-grey dots indicate genes with significant (FDR < 0.05) associations with differentiation. All other dots are colored by the LMM Z-score. (**m**) Left, heatmap of normalized expression levels (rows) for genes associated (LMM FDR < 0.05) with differentiation status (columns). Right, UMAP embeddings illustrating normalized expression levels for representative genes associated with differentiation status.

Given that dedifferentiation characteristics were apparent within six days of culture, we aimed to characterize the transcriptional dynamics associated with culturing diverse protoplasts cell identities. To this end, we generated replicated single-cell RNA sequencing (scRNA-seq) libraries from Columbia-0 Arabidopsis protoplasts cultured for 0, 2, 4, and 6 days in liquid media with and without phytohormone supplementation (**Fig. 1a**; **Table S1**). Following strict quality control thresholds, we identified 125,091 single cells with an average of 2,871 unique molecular identifiers and 1,082 expressed genes per cell, enrichment of nuclear-derived transcripts (>70%), and strong reproducibility among biological replicates (Pearson’s correlation coefficient range = 0.89-0.98; **Fig. S1b**-**S1g**). Clustering and annotation of protoplasts exclusively from Day 0 conditions recovered 21 differentiated cell types stereotypical of Arabidopsis seedlings, including the expected somatic cell identities and cell type distributions (**Fig. 1e**-**1f**; **Fig. S2a**-**S2f**). Integration of all time points and treatments uncovered extensive cell identity reprogramming in protoplasts with extended culture durations and phytohormone applications (**Fig. 1g; Fig. S3a**-**S3e**). For example, we identified seven clusters significantly depleted of somatic (Day 0) cells (Chi-square test, log_2_ fold change < -3, FDR < 0.05) with widespread differential gene expression relative to differentiated cell states (*n*=9,307 genes; per cluster range = 2,509-5,762 genes; FDR < 0.05; **Fig. S3e**-**S3g**; **Table S2**). Investigating fold-change directions revealed that somatic-depleted clusters had a greater proportion of down-regulated genes compared to other clusters, suggesting that somatic identity programs are decommissioned by protoplast culturing (average fraction down-regulated = 0.80 vs. 0.52; Wilcoxon rank sum test, *P-*value < 1.1e^-4^; **Fig. S3h**). Indeed, GO analysis of down-regulated genes in clusters depleted of somatic cells were associated with biological functions such as photosynthesis, primary and secondary metabolism, and morphogenic and developmental processes (FDR < 0.05; **Table S2**). In contrast, up-regulated genes in clusters depleted of somatic cells were enriched for cell wall biosynthesis, cytokinin and auxin signaling, and embryonic meristem specification and development.

To quantify cellular dedifferentiation across the integrated data set, we estimated differentiation scores that reflect expression pattern similarity with either somatic Day 0 cells or a published Arabidopsis cell type atlas^29^ where new transcription was inhibited prior to protoplast isolation (**Fig. 1h; Fig. S3j**). Differentiation scores were highly correlated between the two reference sets (Pearson’s correlation coefficient = 0.72), indicating that somatic cell states remain largely intact despite substantial transcription reprogramming induced by protoplast isolation (**Fig. S3j**). To simplify downstream analysis, we refer to differentiation scores as the similarity with Day 0 somatic cells for the remainder of the text (**Fig. 1h**). Modeling differentiation scores with multiple linear regression indicated that culture duration was a significant predictor, accounting for 42.3% of variance (ANOVA, *P-*value < 2.2e^-16^; **Fig. 1i**). Phytohormone supplementation also impacted differentiation status; hormone-treated cells were more transcriptionally diverged from somatic identities compared to non-treated cells from the same culture duration (Wilcoxon rank sum test, *P-*value < 1.51e^-73^; **Fig. 1i**). Despite evidence of extensive reprogramming, ∼25% of cells from extended culturing (2+ days) maintained transcriptional profiles consistent with Day 0 somatic cell identities (Differentiation score > 0.5; **Fig. S3i**, **Fig. S4a**-**S4b**). However, the proportions of somatic cell identities changed significantly over time (Chi-square test, FDR < 0.05; **Fig. 1j**). While the fraction of dedifferentiated cells (Differentiation score < 0.5) increased regardless of media composition, we found cell composition changes depended on phytohormone treatments (**Fig. 1j**). For example, the proportion of protoplasts with palisade mesophyll-like identity in phytohormone-deficient conditions increased 1.8-fold after four days (Chi-square test, FDR < 2.2e^-16^; **Fig. 1j**). In the presence of phytohormones, the proportion of meristem-like cells increased more than 18-fold by day six (Chi-square test, FDR < 2.2e^-16^; **Fig. 1j**). In fact, cells with protoderm- and meristem-like identity were the only over-represented somatic cell types after six days of culturing with phytohormones, mapping predominantly to clusters we identified as being depleted of Day 0 somatic cells (**Fig. 1g**, **1j**, **1k**, **Fig. S4a**-**S4b**). These results suggest that phytohormone treatments either help maintain existing meristematic cell states in vitro or stimulate transcriptional reprogramming of somatic cell types towards meristem-like states.

Quantification of somatic cell identity provides an opportunity to pinpoint transcriptional signatures underlying dedifferentiation gradients. To this end, we assessed the relationship between patterns of gene expression and differentiation scores across cells using linear mixed effects models (LMM) to control for pseudoreplication bias and other technical covariates (**Fig. 1l**-**1m**). In total, we uncovered 2,740 and 9,921 genes with significant positive and negative effects on differentiation status, respectively (FDR < 0.05; **Fig. 1l**-**1m**; **Table S2**). The top associated genes were consistent with salient signatures of somatic functions. For example, expression of photosynthesis genes was strongly associated with differentiation (**Fig. 1l**-**1m**). In contrast, dedifferentiation was associated with expression of cytokinin/auxin response genes, chromatin remodelers (*SPLAYED; SYD*), and TFs previously associated with wound responses, including *AP2/ERF* family TFs *WIND1*, *ERF115*, and *RELATED TO APETALA-2.6 LIKE* (*RAP2.6L*; **Fig. 1m**, **Fig. S4c**). We also observed expression of known shoot meristem regulators, such as *ESR1*^8^, specifically expressed in meristem-like cells (cluster 7) unique to phytohormone treated samples (**Fig. 1m**). Gene set enrichment analysis (GSEA) of differentiation gradient-associated genes revealed terms related to chloroplast biogenesis, photosynthesis, primary metabolism, and shoot/leaf development (FDR < 0.05; **Table S3**). In contrast, dedifferentiation-associated genes were enriched for terms related to embryonic meristem development, meristem initiation and specification, meristem structural organization, response to wounding, and auxin biosynthesis. Collectively, these analyses highlight the principal transcriptional and cell state dynamics along the general axis of dedifferentiation.

### Phytohormone treatment decelerates protoplast aging

We hypothesized that modeling cell age as a function of transcriptional state and hormone treatment would be informative for understanding the effects of auxin and cytokinin on protoplast reprogramming. To test this, we split cells by hormone treatment status for matched culture times (Day 0 to Day 4) and fit elastic net models to learn the predictive transcriptomic signatures of protoplast age for each treatment independently (**Fig. 2a**). Comparison of predicted and ground truth cell age from held-out data revealed that the elastic net models robustly capture a continuous representation of cell age (real time cell age) based solely on transcriptomic information (Pearson’s correlation coefficient [PCC] > 0.95; **Fig. 2b**). Next, we made predictions for all combinations of transcriptomic inputs and trained models, for example, predicting real time age of non-hormone treated cells using the model trained on hormone-treated cells. We then calculated the difference between predicted cell age and ground truth (**Fig. 2c**). Both models were able to accurately predict the age of nascent (Day 0) protoplasts regardless of the treatment status of withheld test cells. However, we observed that the model trained on non-hormone treated samples significantly underestimated the age of hormone-treated cells, while the model trained on hormone-treated cells overestimated the age of non-hormone treated cells, suggesting that hormones affect the transcriptomic features that are predictive of cell age (Wilcoxon signed-rank test, *P-*value < 2.2e^-16^; **Fig. 2c**). Investigating further, we found expression of genes with aging and senescence ontologies were delayed in hormone-treated cells, whereas signatures of cell proliferation were enriched (**Fig. S5a**-**S5b**). These results suggest a relationship between phytohormone-stimulated cell proliferation and attenuation of protoplast aging.

**Figure 2:**
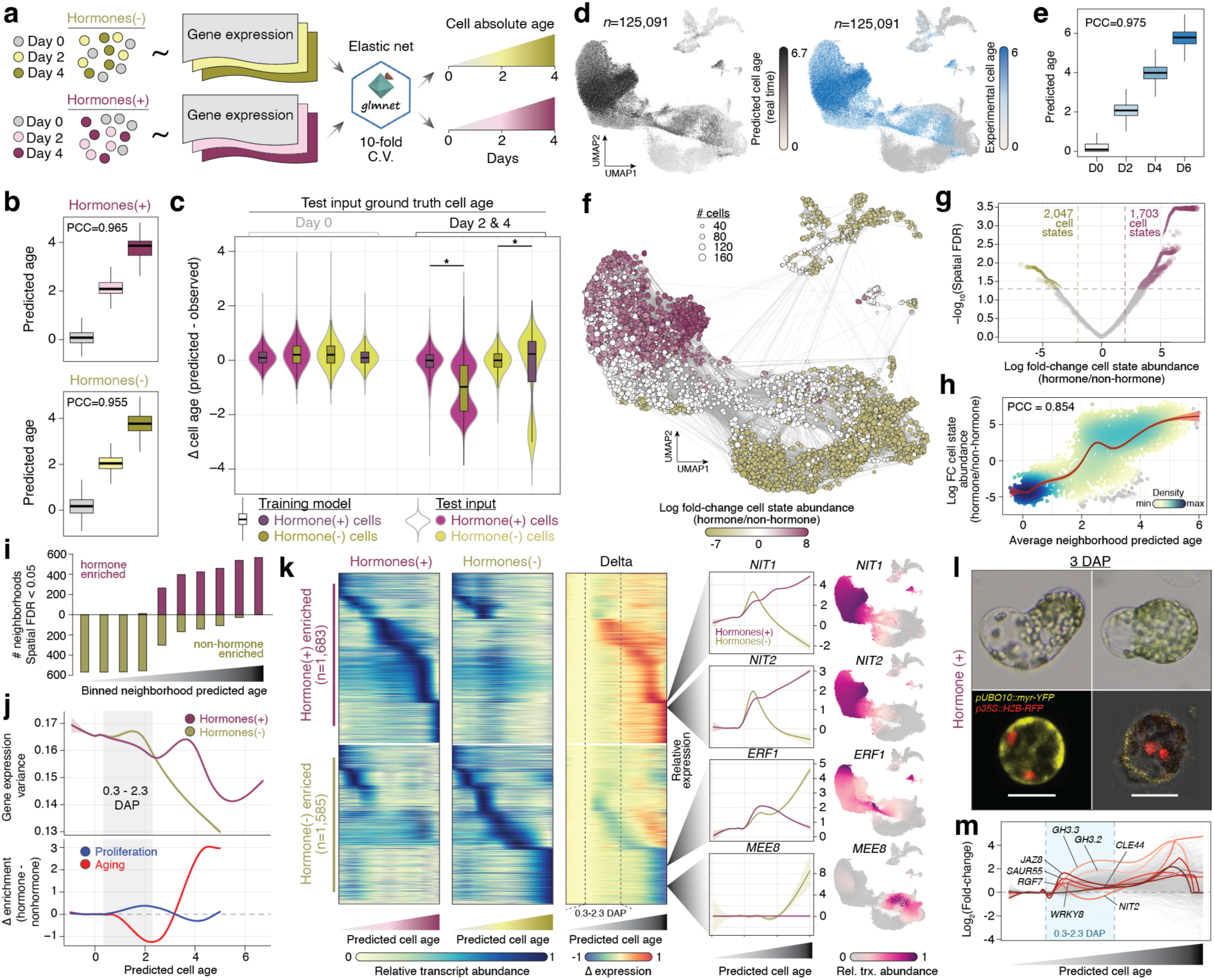
Phytohormone application attenuates protoplast aging and promotes cell proliferation. (**a**) Schematic of elastic net training and prediction strategy. (**b**) Distribution of predicted cell ages for each ground truth classification for models trained with hormone-treated cells (top) and non-hormone-treated cells (bottom) using time-matched samples. (**c**) Distributions of cell age delta between predicted and ground truth cell ages for each combination of training model (hormone and non-hormone-trained model are distinguished by boxplot color) and input protoplast transcriptomes (hormone and non-hormone single cell transcriptomes are distinguished by violin color) used to make predictions. (**d**) UMAP embeddings of predicted cell age in the full data set from hormone-treated and non-hormone-treated elastic net models (left) and experimentally measured cell age (right). (**e**) Distributions of predicted cell ages for each ground truth protoplast age classification from the fully specified models. (**f**) UMAP embedding of neighborhoods colored by log_2_ fold-change enrichment of hormone over non-hormone cell states after accounting for total cell counts per treatment. Grey lines indicate connected neighborhoods. (**g**) Volcano plot of differential neighborhood enrichment. (**h**) Density scatter plot of average neighborhood predicted age (x-axis) versus log_2_ fold-change enrichment. Shaded regions around the fitted line indicates the 95% confidence interval. (**i**) Number of significant neighborhoods enriched with non-hormone (gold) and hormone (purple) cells across 10 equal sized bins partitioned by average predicted age. (**j**) Top, gene expression variance across predicted cell age. Shaded regions around fitted lines indicate 95% confidence intervals. Bottom, delta (hormone – non-hormone) enrichment for aging (red) and proliferation (blue) gene sets. (**k**) Heatmaps of temporal expression values for hormone, non-hormone, and the delta between hormone and non-hormone expression levels across real time (left). Exemplary differentially expressed genes plotted across real time and on the UMAP embedding (right). Shaded regions around fitted lines indicate 95% confidence intervals. (**l**) Top; representative protoplast morphologies at 3 days after protoplast (DAP) induction in the hormone treated cultures. Bottom; confocal micrographs of dividing 3 DAP with labeled cell membranes (*pUBQ10::myr-YFP*) and nuclei (*p35S::H2B-RFP*). Scale bar = 10 μm. (**m**) Differential expression (log_2_ fold-change) between hormone and non-hormone-treated cells across predicted cell age. Exemplary early onset differentially expressed genes are colored in a red hue. All other differentially expressed genes are colored in grey.

Next, we compared the genes with predictive power between the two models. Unexpectedly, coefficient estimates between the two models were uncorrelated (PCC = -0.016), indicating that the transcriptional signatures predictive of cell age are distinct between phytohormone regimes (**Fig. S5c**). To investigate this, we applied fully specified elastic net models for hormone- and non-hormone-treated cells to predict continuous real time cell ages in the full data set (**Fig. 2d**). As with the reduced data set, the fully specified models accurately captured experimental age (PCC = 0.975), supporting the suitability of transcriptome data for modeling age phenotypes (**Fig. 2e**). Next, we quantified the effect of phytohormone applications on cell state heterogeneity by defining unique transcriptomic states across the real time predictions using a neighborhood-based framework (**Fig. 2f**-**2g**). This analysis revealed that phytohormone treatments were associated with greater cell state diversity with increasing real time, suggesting exogenous auxin and cytokinin application promotes transcriptional variation (Spatial FDR < 0.05; **Fig. 2h**-**2i**). Although gene expression variance generally decreased over real time regardless of treatment, consistent with dedifferentiation of somatic cell types with extended protoplast culturing, we observed significantly greater gene expression variance in phytohormone treated cells beginning at ∼2.3 days after protoplast isolation (DAP; Likelihood ratio test, *P* < 2.2e^-16^; **Fig. 2j**). The onset of enhanced transcriptional diversity in phytohormone-treated samples coincided with the timing of cell aging attenuation and cell proliferation gene program enrichment (**Fig. 2j**).

To understand the effect of culture duration and treatment on transcriptional reprogramming, we first identified genes with dynamic expression across real time. Of the 27,841 expressed genes, 53.3% (*n*=14,848) vary significantly as a function of real time (FDR < 0.05; **Fig. S5d**). We hypothesized that hormone treatment could influence time-dependent regulation. To test this, we characterized differential gene expression between age-matched hormone and non-hormone-treated cells across real time using generalized additive mixed models. We found a total of 3,268 genes with temporally dynamic expression with varying patterns between hormone treatments (FDR < 0.05; **Fig. 2k**). Of these, 1,683 and 1,585 genes were upregulated in hormone and non-hormone treatments, respectively. The top hormone-biased and time-dependent genes included many known auxin biosynthesis genes, while non-hormone treated cells were associated with genes related to embryogenesis and regulation of stress response (**Fig. 2k**). GSEA revealed terms such as “hormone regulation”, “cell wall assembly”, and “cell-cell junction” enriched in genes associated with temporal hormone signaling, consistent with prevalent nuclear replication phenotypes in 3 day-old cell membrane (*pUBQ10::myr-YFP*) and nuclei-tagged (*p35S::H2B-RFP*) protoplasts in hormone treated cultures (**Fig. 2l**; **Table S4**). Conversely, non-hormone treated cells were associated with terms related to photosynthesis, response to light, and ribosome biogenesis, highlighting distinct regulatory trajectories stimulated by phytohormone perturbation (**Table S4**).

Next, we aimed to pinpoint genes associated with the onset of increased transcriptional variation at ∼2.3 DAP in the phytohormone-treated cultures. A total of 438 genes were differentially expressed between treatments within a 2-day window preceding 2.3 DAP (0.3-2.3 DAP; **Fig. 2k**; **Fig. S5e**). As expected, ranking genes by the timing of expression divergence between treatments revealed early activation of many known auxin-response genes, including *GRETCHEN HAGEN 3.3/3.2, NIT2,* and *SMALL AUXIN UPREGULATED RNA 55* (*SAUR55*; **Fig. 2m**). We also identified other known hormone signaling pathways induced by auxin and cytokinin treatment (**Fig. 2m**). For example, *JASMONATE ZIM-DOMAIN PROTEIN 8* (*JAZ8*) was among the first differentially expressed genes following hormone treatments. At basal levels of jasmonates (JA), JAZ8 recruits the corepressor TOPLESS to maintain transcriptional repression of JA-responsive transcription factors^30^. Activation of *JAZ8* suggests auxin/cytokinin signaling decommissions JA-induced wounding responses and helps to direct transcriptional reprogramming towards reestablishing multicellularity, consistent with previously established interplay between JA and auxin^31^. Indeed, *MYC2/3/4*, master regulators of JA transcriptional responses, including cross and autoregulation, and known targets of JAZ8^30,32^, all exhibited downregulation within the 2-day window following the onset of *JAZ8* upregulation (**Fig. S5f**). Consistent with reprogramming towards stem cell identity, hormone-biased *JAZ8* expression was concomitant with increased expression of *ROOT MERISTEM GROWTH FACTOR 7*, a positive regulator of *PLT* family genes and meristem identity^33^, followed by markers of stem cell identity, such as *CLAVATA3/ESR-RELATED 44* (*CLE44*; **Fig. 2m**)^34^. Taken together, these data indicate that phytohormones promote large-scale transcriptional reprogramming, repression of master regulators underlying wounding responses, and activation of genes implicated in stem cell fate.

### Dedifferentiation potential of diverse somatic cell identities

Dedifferentiation potential of diverse somatic cell types remains poorly understood. To address this, we first identified putative dedifferentiated cell populations by manually evaluating expression of stem cell marker genes, dedifferentiation scores, treatment and culture duration, cluster relatedness, and retention of somatic cell identities (**Fig. 3a**; **Fig. S5g**). This analysis nominated five distinct non-somatic cell populations with low diffusion component values, an independent estimate of differentiation status, that emerged under varying treatment conditions: cluster 4, enriched for non-hormone-treated protoplasts, and clusters 6, 7, 18, and 21 associated with hormone-treated cells (**Fig. 3b**). Unexpectedly, we found that expression of stem cell marker genes, such as *LEC2*, *PLT7, PHAVOLUTA* (*PHV*)*, LBD16*, and *MONOPTEROS* (*MP*) varied widely across dedifferentiated clusters (**Fig. 3a**-**3b**; **Fig. S5h**). Application of GSEA revealed clusters 4, 6, 18, and 21 were enriched for terms related to embryogenesis and the cell cycle, and only cluster 6 was associated with “embryonic meristem development, initiation, and structural organization” (**Fig. S6**; **Table S5**). Indicative of heterogenous reprogramming, cluster 7 was associated with GO terms related to “tissue morphogenesis” and expression of genes associated with shoot regeneration (*ESR1*), cell wall assembly (*EXTENSIN 3*), root meristem regulation (*LBD11*), and callus/primordia development (*LCR69, PDF2.1, GRP4, FIB2;* **Fig. 1k**, **Fig. 3a**, **Fig. S5h**). We observed co-expression of well-known embryogenic markers *LEC1, YUC4*, *SOMATIC EMBRYOGENESIS RECEPTOR KINASE 1* (*SERK1*), *AGL15*, and *BBM* in protoplasts with S-phase and remnant mesophyll-like identity (cluster 21), while *LEC2, AGL15,* and *WIND1* were enriched within a dedifferentiated cluster that retained a notable fraction of somatic guard cells (cluster 18; **Fig. 3a**, **Fig. S5h**). Curiously, the unexpected expression of *LEC1/2* in mesophyll and guard-like cells, respectively, was observed immediately following protoplast generation less than 4 hours from intact tissue (**Fig. 3a**, **Fig. S5h**). However, we found that most protoplasts were hallmarked by *PLT7*, *PHV*, *MP* and *ABI3* expression, consistent with the majority of dedifferentiating cells following a root-associated developmental pathway (**Fig. 3a, Fig. S5h**)^23^. Importantly, the activation of stem cell markers was unique to protoplasts; stem cell markers were uniquely enriched in protoplasts compared to tissue-matched single-nuclei RNA-seq data, and in contrast to other expressed genes (Wilcoxon rank sum, *P*<2.6e^-3^; **Fig. S5i**-**S5k**). Although less than 1% of cultured protoplasts ultimately give rise to callus, these data indicate that cell wall digestion induces expression of diverse stem cell marker genes across most cultured cells in an manner that depends on initial somatic cell identity.

**Figure 3:**
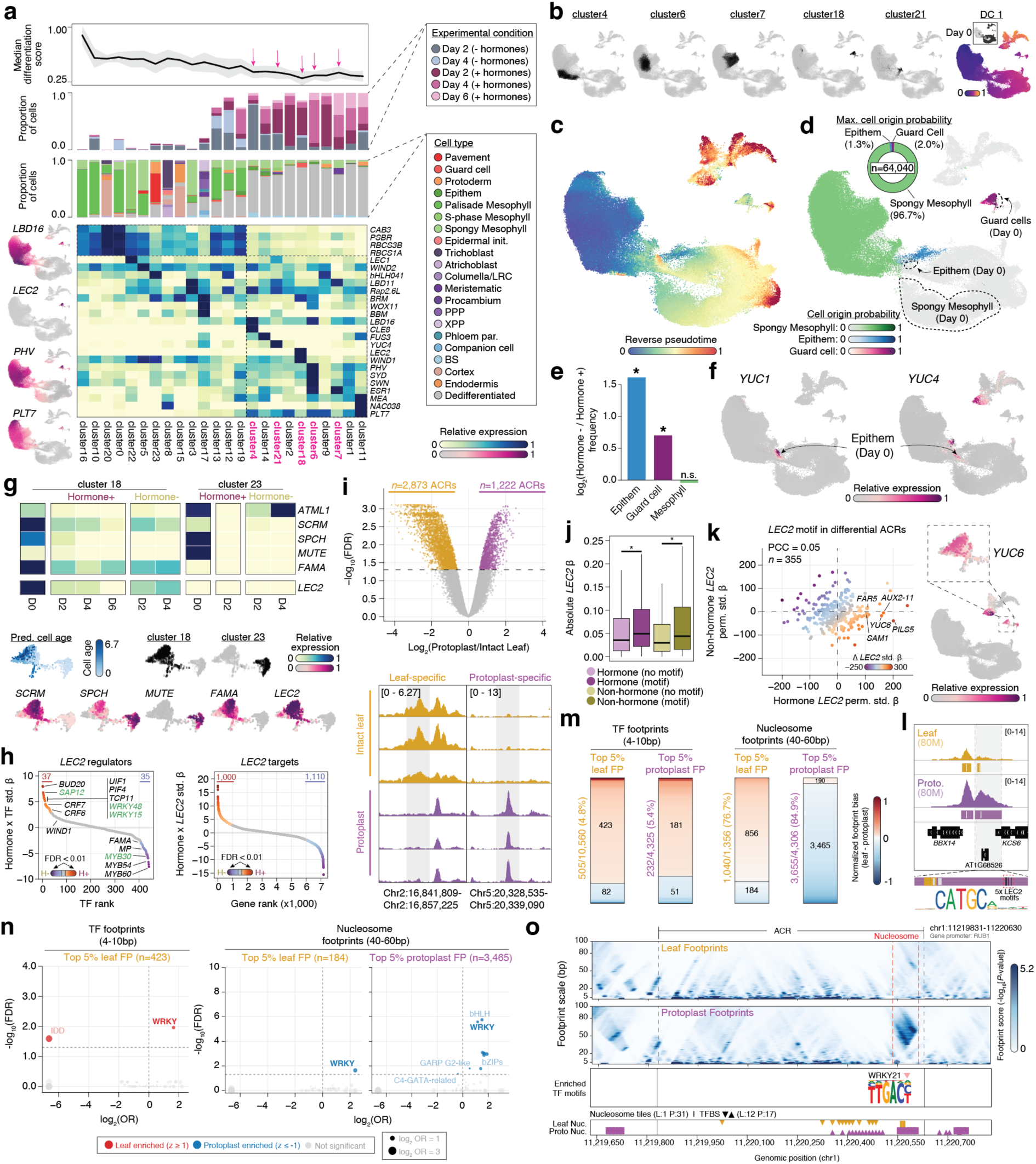
Dedifferentiation potential is biased by somatic cell identity. (**a**) Annotated heat map illustrating row-normalized expression levels of marker genes across clusters. Top panel, line graph illustrating the median (black line) and standard deviation of differentiation scores within each cluster. Middle panel, proportion of cells partitioned by experimental condition. Bottom panel, proportion of cells partitioned by cell type. Exemplary normalized expression levels for four marker genes across cells on the UMAP embedding are shown to the left of the heatmap. (**b**) UMAP visualization of cells from cluster 4, 6, 7, 18, and 21 (black dots). Diffusion component 1 values are illustrated in the UMAP on the right. The inset illustrates Day 0 somatic cells (black dots). (**c**) UMAP embedding of the integrated data set colored by the consensus reverse pseudotime estimates. (**d**) UMAP embedding illustrating cell origin probabilities for the 10 dedifferentiated cell clusters. Inset, donut chart showcasing the proportion of cells with maximum cell origin probabilities for spongy mesophyll, guard cell, and epithem cell identities. Day 0 cells corresponding to spongy mesophyll, epithem, and guard cells are outlined with black dashed lines. (**e**) Log_2_ transformed ratio of non-hormone to hormone-treated maximum cell origin frequency. Asterisks denote *P*<1.1e^-15^. (**f**) UMAP embedding illustrating expression levels of *YUC1* and *YUC4*. The centroid of Day 0 epithem cells is denoted by the black arrows. (**g**) Heatmap visualizing expression levels of stomatal lineage and epidermal TFs along with *LEC2* partitioned by cluster and hormone treatment status. UMAP embedding of cluster 18 and 23 cells and expression of exemplary stomatal lineage TFs and *LEC2* are shown below. (**h**) Left; ranked standardized effects of the interaction between hormone treatment and time-lagged TF expression on *LEC2* regulation. Positive interaction terms represent TF regulators of *LEC2* where hormone treatment promotes TF contribution to *LEC2* expression. Right; ranked standardized effects of the interaction between hormone treatment and *LEC2* expression on *LEC2* target genes. Colored dots represent significant (FDR < 0.01) interactions. (**i**) Top; volcano plot of differentially accessible chromatin regions (FDR < 0.05) between protoplast (purple) and intact leaves (orange). Bottom; exemplary differentially accessible chromatin regions associated with intact leaves (orange) and protoplasts (purple). (**j**) Distribution of absolute *LEC2* effects on target genes near ACRs with (dark) and without (light) LEC2 motifs partitioned by hormone (purple) and non-hormone (yellow) time-lagged gene regulatory networks. (**k**) Left; scatterplot illustrating hormone (x-axis) and non-hormone (y-axis) permutation standardized *LEC2* effects on *LEC2* target genes near differential ACRs with LEC2 motifs. Right; UMAP embedding illustrating relative transcript abundance of *YUC6.* Inset, cluster 18 cells with guard-like identity. (**l**) Genome browser of an exemplary ACR expanded in protoplast samples containing a cluster of 5x LEC motifs. ACR coordinates are demarcated by the orange (intact leaf) and purple (protoplast) bars. ACR summits are depicted with grey vertical lines. Black bars represent LEC2 motif occurrences. (**m**) Distribution of leaf and protoplast-biased TF and nucleosome-scale footprints in leaf-specific and protoplast-specific ACRs. (**n**) TF family motif enrichment within TF- and nucleosome-scale footprints for leaf- and protoplast-specific ACRs. (**o**) Multiscale footprinting example at the *RUB1* promoter.

To establish the dedifferentiation potential of distinct somatic cells under our experimental conditions, we modeled reverse pseudotime trajectories towards all dedifferentiated cell clusters, defined as clusters composed of >50% dedifferentiated cells (*n* clusters = 10), using probabilistic cell fate diffusion mapping (**Fig. S7a**). Evaluation of the cell fate maps revealed canonical developmental trajectories, such as specification of atrichoblast, trichoblast, and lateral root cap from root epidermal initials, bifurcation of cortex and endodermal cell lineages, and above-ground epidermal cell types derived from the meristematic protoderm (**Fig. S7b**). To simplify downstream analysis, we defined the consensus reverse pseudotime per cell as the minimum reverse pseudotime across all 10 fitted trajectories (**Fig. 3c**; **Fig. S7a**). Consensus reverse pseudotime estimates were strongly supported by several orthogonal metrics, including differentiation potential based on transcriptional diversity (CytoTRACE)^35^, real time cell age predictions, and the initial correlation-based differentiation scores (**Fig. 1f**; **Fig. 2d**; **Fig S8a**-**S8d**). Next, we reasoned that somatic cell fate probabilities towards dedifferentiated clusters could be interpreted as cell origin probabilities. Classifying the 10 dedifferentiated clusters (*n* cells = 64,040) by the most probable somatic origin revealed a reduced set of above ground cell types – mesophyll, epithem, and guard cell-like identities – as the major source of dedifferentiated cells (**Fig. 3d**; **Fig. S7b**, **Fig. S8e**). The proportion of dedifferentiated cells derived from mesophyll (97%) and guard cells (2%) was nearly identical to a prior study of 164 alginate-embedded protoplasts that underwent cell division following phytohormone treatments (98.78% mesophyll-like, 1.22% guard cells; **Fig. 3d**)^15^. Unlike somatic mesophyll and guard cells, dedifferentiated cells with epithem origin has not been previously reported. Further investigation indicated that dedifferentiated cells derived from epithem and guard cell-like identities were significantly enriched in the non-hormone treated cultures (Chi-square test; *P <* 1.11e^-15^; **Fig. 3e**). Epithem cells comprising hydathodes are centered on auxin maxima at the margins of Arabidopsis leaves^36^, leading us to speculate that epithem dedifferentiation may be promoted by cell autonomous auxin biosynthesis. We observed elevated expression of auxin biosynthesis genes, such as *YUC1* and *YUC4*, response to auxin stimulus, and regulation of auxin polar transport program within somatic epithem cells (**Fig. 3f**; **Fig. S8f**). Hormone treatments enhanced expression of auxin biosynthesis genes between 0 and 2.5 days, potentially producing lethal levels of auxin, in contrast to non-hormone treated protoplasts (**Fig. S8g**). Furthermore, reduced levels of auxin biosynthetic gene expression in non-hormone cells was associated with a greater rate of dedifferentiation (**Fig. S8h**). Collectively, these results indicate that dedifferentiation potential depends on the interaction between somatic cell identity and auxin-related processes.

### Ectopic *LEC2* activity is influenced by phytohormone treatments in guard cells

Analysis of stem cell marker genes indicated the unexpected expression of several genes involved in embryogenesis, including *LEC2*^37^. To better understand the role of *LEC2* in dedifferentiation, we queried the dynamic gene regulatory networks and cell states associated with its expression. *LEC2* transcripts were primarily produced in protoplasts with guard cell identity, hallmarked by *WIND1*, *SCREAM, SPEECHLESS*, and *FAMA* expression (**Fig. 3g**; **Fig. S5h**). *LEC2* expression was greatest at Day 0 and decreased over time in phytohormone-treated protoplasts, while *LEC2* expression increased between Day 2 and 4 in non-hormone cultures, indicating that exogenous phytohormones antagonizes *LEC2* transcription. To investigate time- and hormone-dependent factors associated with *LEC2* expression, we constructed time-lagged gene regulatory networks for hormone and non-hormone treated dedifferentiating guard cells using regularized ridge regression (permutation FDR < 0.01; **Fig. S9a**-**S9b**; **Table S6**-**S7**). LEC2 target gene effects in both hormone and non-hormone treated cells were similarly associated with *LEC2*-induced expression in prior experiments, corroborating the inferred networks (**Fig. S9c**-**S9d**)^38^. However, comparison of hormone and non-hormone networks revealed moderate correlations for TF effects on *LEC2* transcript abundance (PCC = 0.5), but not for LEC2 targets (PCC = 0.08; **Fig. S9a**-**S9b**), suggesting hormone effects are downstream of *LEC2* activation.

To investigate the impact of phytohormones in more detail, we generated a conditional time-lagged gene regulatory network and assessed interactions between TF effects and hormone treatments using permutation-based testing. This analysis revealed 72 putative *LEC2* regulators and 2,110 putative *LEC2* targets with significantly different time-lagged effects between hormone and non-hormone treated protoplasts (FDR < 0.01; **Fig. 3h**; **Table S8**). The proportion of LEC2 targets (29%) affected by hormone treatments was nearly twice the proportion of *LEC2* regulators (16%), consistent with phytohormones primarily affecting LEC2 downstream activity (Chi-squared test; *P-*value *<* 5.1e^-6^). Expectedly, several TFs implicated in cytokinin and auxin signaling, such as *CYTOKININ RESPONSE FACTOR 6/7* and *MP* were associated with significantly different effects on *LEC2* transcript levels depending on treatment (**Fig. 3h**). TFs associated with upregulation of *LEC2* in the phytohormone treatment were enriched for GO terms related “maintenance of meristem identity”, “stem cell development”, and “stem cell maintenance” (*P-*value < 4.9e^-3^), while guard identity genes, *FAMA* and *MYB60,* and several stress responsive TFs were associated with *LEC2* expression in non-hormone treatments (**Fig. 3h**). These results highlight complex interplay between hormone signaling, stress responses, and cell identity as important contributors to *LEC2* activation and activity in dedifferentiating guard cells.

### Dedifferentiation induces widespread regulatory rewiring

We posited that *LEC2* activity in protoplasts could be captured by changes in chromatin accessibility at LEC2 targets. Using existing ATAC-seq data derived from nascent protoplasts and intact leaves, we identified 4,095 differentially accessible chromatin regions (ACRs; FDR < 0.05; **Fig. 3i**)^16^. Compared to all ACRs (15.8%), differential ACRs between intact leaves and protoplasts (22.0%) were enriched for LEC2 motifs (Chi-squared test; *P*-value < 2.7e^-15^). Reciprocally, ACRs with LEC2 binding sites were associated with significantly greater chromatin accessibility variability compared to ACRs lacking LEC2 motifs (Wilcoxon rank sum test; *P-*value < 7.6e^-21^; **Fig. S9e**). Time-lagged *LEC2* effects on genes (*n*=355) near differentially accessible LEC2 binding sites were significantly greater compared to those lacking LEC2 binding sites in both hormone and non-hormone treated samples, suggestive of LEC2 targeting (Wilcoxon Rank Sum Test, *P-*value < 7.8e^-7^; **Fig. 3j**). Putative targets with accessible LEC2 motifs included several auxin-related and lipid metabolism genes, such as *YUC6*, *AUXIN INDUCIBLE 2-11, PIN-LIKES 5, FATTY ACID REDUCTASE 5,* and *S-ADENOSYLMETHIONINE SYNTHETASE 1,* reminiscent of the canonical function of LEC2 in embryogenesis (**Fig. 3k**)^26^. *LEC2* effects on putative target genes were largely distinct between hormone and non-hormone-treated protoplasts (PCC = 0.05, concordance = 44.7%).

Distinct LEC2 effects on targets between treatments could arise from cooperative regulation. Indeed, differential ACRs with LEC2 motifs were significantly larger, contained more diverse TF binding sites, and were associated with significant expansion following protoplast isolation compared to differential ACRs lacking LEC2 motifs (Wilcoxon rank sum test; *P-*value < 2.1e^-20^;**Fig. S9f**-**S9h**). For example, we identified an expanded ACR downstream of *3-KETOACYL-COA SYNTHASE* (*KCS6*), part of the long chain fatty acid synthesis pathway, containing a cluster of five LEC2 binding sites uniquely accessible in protoplasts (**Fig. 3l**). We posited that TFs differentially induced by hormone treatment could act on these expanded ACR domains, leading to hormone-specific regulatory outcomes. To test this, we estimated TF co-motif enrichment in LEC2-associated ACRs that gained or lost chromatin accessibility upon protoplast isolation relative to stable ACRs. Protoplast-specific ACRs with LEC2 motifs were strongly enriched for ABI3, FUS3, and several homeodomain TF binding sites, consistent with the LEC2-ABI3-FUS2 embryogenic module (FDR < 0.05; **Fig. S9i**-**S9j**). Notably, TFs co-enriched with LEC2 had significantly greater phytohormone-dependent effects on LEC2 targets compared to non-target genes (Wilcoxon rank sum test; *P*<2.2e^-3^; **Fig. S9k**) and were strongly differentially expressed between hormone treatments at the onset of in vitro culture (FDR < 0.05; **Fig. S9l**). These results suggest that phytohormones promote early differential expression of TFs capable of targeting expanded accessible chromatin near LEC2 target genes.

Cell wall digestion also induced substantial loss of somatic chromatin accessibility signatures. Somatic ACRs condensed in protoplasts were strongly enriched for motifs recognized by WRKY family TFs (FDR < 0.05; **Fig. S9i**). The coincidence of condensed WRKY binding sites was at odds with *WRKY* expression dynamics after protoplast induction; 52% (23/44) of *WRKY* TFs exhibit similar or increased expression, while the motifs of all (44/44) WRKY TFs were enriched in newly compacted chromatin (**Fig. S9m**). In fact, TF binding sites for several upregulated and highly expressed *WRKY* TFs, such as *WRKY28* and *WRKY6*, were among the most enriched in condensed chromatin unique to protoplasts (**Fig. S9n**). The association between WRKY binding sites and chromatin compaction was not limited to ACRs with LEC2 binding sites but generalized to the entire genome (**Fig. S9o**-**S9p**). The most upregulated *WKRY* TFs from bulk RNA-seq (*WRKY6/16/28*) were also strongly expressed in both nascent and extended protoplasts cultures, with expression levels greatest at the transition between 0- and 2-day old protoplasts (**Fig. S9q**). The motifs of the top upregulated and downregulated *WRKY* TFs were similar, suggesting that protoplast-induced *WRKY* TFs may compete or displace somatic *WRKY* TFs (**Fig. S9r**). To investigate this further, we analyzed dynamic protein-DNA architectures within protoplast-condensed ACRs via multiscale footprinting (**Fig. S10a**-**S10m**)^39^. We observed a systematic decrease in TF-scale (4-10bp) footprint scores in ACRs losing chromatin accessibility in protoplasts, while nucleosome-scale (40-60bp) footprints were strongly enriched, consistent with destabilized TF binding and increased nucleosome occupancy (**Fig. 3m**). Analysis of TF motifs revealed that sequences recognized by WRKY TFs were associated with a transition from leaf-biased TF-scale footprints to protoplast-biased nucleosome-scale occupancy (**Fig. 3n**). For example, we identified a TF-scale WRKY footprint enriched in leaf tissue in the *RELATED TO UBIQUITIN 1* (*RUB1*) promoter that was replaced by a nucleosome-scale footprint in protoplast chromatin (**Fig. 3o**). Collectively, these data suggest that protoplast-induced WRKY TFs decommission a subset of somatic ACRs by enhancing nucleosome occupancy at potentially competitive WRKY binding sites.

### Early regulators of dedifferentiation are independent of somatic cell identity

How initial somatic identity and phytohormones interact across time to dictate dedifferentiation outcomes remains unclear. To address this, we compared mesophyll, epithem, and guard cell dedifferentiation trajectories using 1,000 probabilistic walks from terminal somatic cells to dedifferentiated clusters constrained by decreasing pseudotime and cell-cell distance within the multi-scale reduced dimensional space (**Fig. 4a**; **Fig. S11a**). Probabilistic walks were performed on the full integrated dataset agnostic to experimental conditions, allowing us to ask if phytohormones affect dedifferentiation outcomes. Logistic regression within a generalized additive modeling framework indicated that most probabilistic walks (54%) were associated with a significant increase in the proportion of hormone-treated cells with increasing dedifferentiation across trajectories (FDR < 0.05; **Fig. 4b**). Phytohormone-biased dedifferentiation was most commonly observed for mesophyll trajectories (7/9), whereas guard cell and epithem probabilistic walks were largely unassociated with phytohormones (**Fig. 4b**). Prevalent phytohormone effects on dedifferentiation suggests that hormone treatments may influence the rate of dedifferentiation. To test this hypothesis explicitly, we estimated dedifferentiation velocity – reflecting the rate of change from somatic to dedifferentiated states – across all mesophyll, guard cell, and epithem trajectories partitioned by hormone treatments as a function of real time. We found that dedifferentiation velocity was greatest between day 0 and day 2 timepoints for mesophyll cells, while guard and epithem cells exhibited steady dedifferentiation velocity across real time (**Fig. 4c**). Although subtle, phytohormone treatments in mesophyll generally enhanced dedifferentiation velocity across trajectories after two days of culturing, coinciding with increased transcription variation associated with exogenous auxin and cytokinin applications (**Fig. 4c**, **Fig. 2j**-**2k**). These data suggest that phytohormones accelerate dedifferentiation and promote cell state diversity in somatic mesophyll cells at a timing that coincides with greater transcriptional variance.

**Figure 4:**
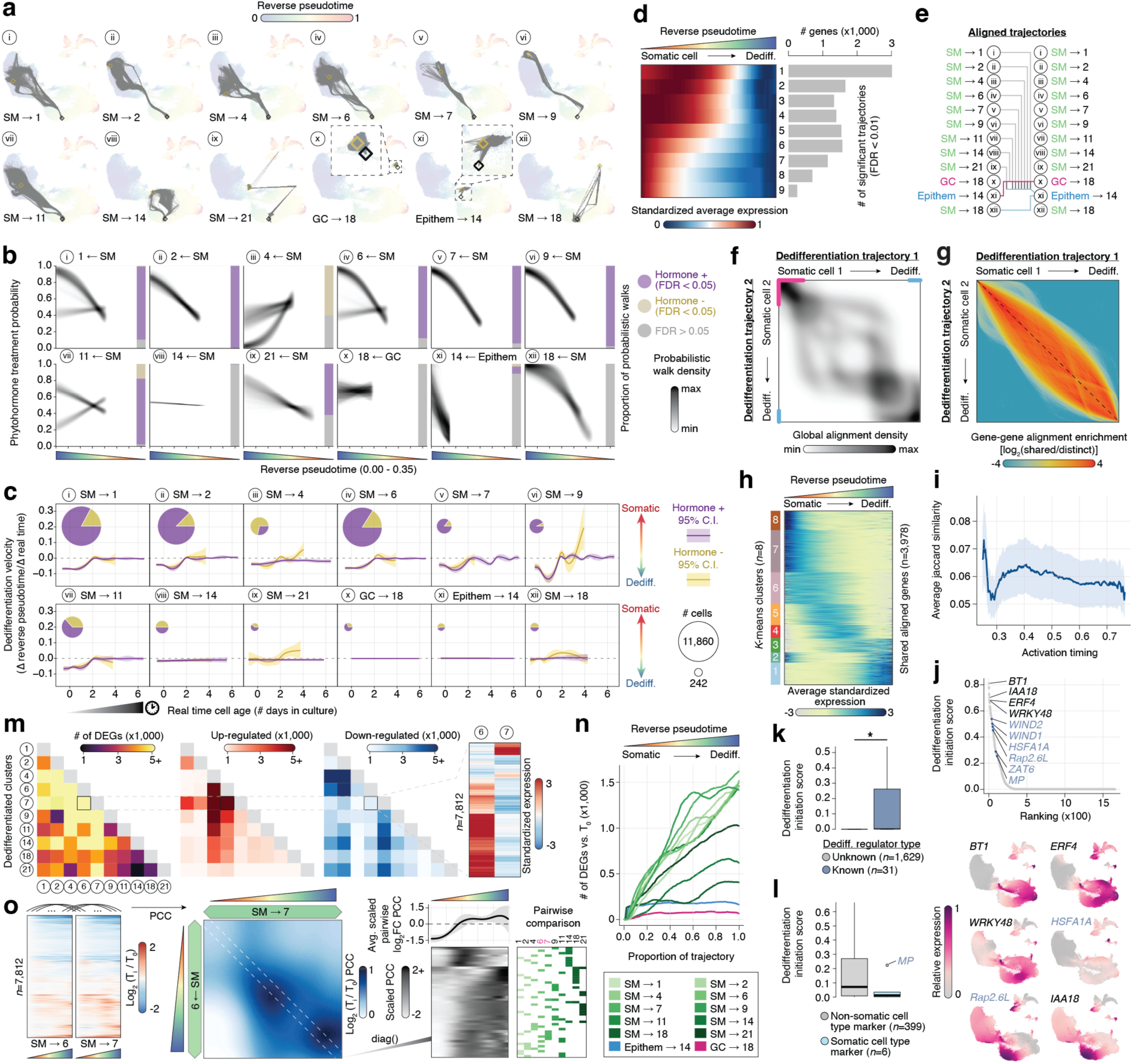
Transcriptional signatures of cell-specific dedifferentiation and cell fate diversity. (**a**) Probabilistic walks (*n*=1,000) from a given somatic cell state (black diamond) to a dedifferentiated cell cluster (gold diamond). SM, spongy mesophyll; GC, guard cell. (**b**) Density scatterplot of logistic regression fits between reverse pseudotime (x-axis) and proportion of hormone-treated cells (y-axis) for each trajectory. Fit intensity reflects the density of probabilistic walks (*n*=1,000). Embedded bars indicate the proportion of probabilistic walks significantly associated (FDR < 0.05) with increasing proportion of hormone-treated cells (purple), non-hormone treated cells (yellow), and non-significant (FDR > 0.05) fits (grey). (**c**) Average and 95% confidence interval of dedifferentiation velocity (rate of pseudotime change per unit time) across time for all 12 dedifferentiation trajectories partitioned by hormone treatment (purple) and non-hormone treated cells (yellow). The total number and proportion of hormone (purple) and non-hormone (yellow) cells are indicated by the pie charts. (**d**) Left, scaled (0-1) averaged gene expression patterns across trajectories partitioned by the frequency of overlap among trajectories. Right, number of genes classified by frequency of overlap among reverse pseudotime probabilistic trajectories. (**e**) Alignment scheme of trajectories with distinct somatic cell identity origins and dedifferentiated cell fates. (**f**) Density heatmap of pairwise dynamic time-warping global gene expression alignments among unrelated dedifferentiation trajectories (pairwise alignments *n* = 19). Pink and blue lines indicates the shared early and late dedifferentiation segments. (**g**) Density heatmap of pairwise gene-gene alignment enrichment (log_2_ shared/distinct) from dynamic time-warping of unrelated reverse pseudotime trajectories. Shared alignments were identified by empirical false discovery control relative to permuted trajectories (eFDR < 0.05). (**h**) Heatmap of gene expression dynamics from somatic to stem-like identity, averaged across trajectories, and then standardized. Colored blocks to the left represent k-means assigned clusters. (**i**) Average Jaccard similarity of regulator target genes within discrete activation timing bins (n=300). The light shaded region represents the standard deviation in Jaccard similarity scores. (**j**) Top, scatter plot representing dedifferentiation initiation score regulator gene rankings. Red dots indicate genes previously implicated in dedifferentiation regulation. Bottom, exemplary UMAP embeddings colored by relative gene expression for select regulators. (**k**) Boxplot of dedifferentiation initiation scores for previously known and unknown dedifferentiation regulators. Outliers have been removed for readability. (**l**) Distribution of dedifferentiation initiation scores for genes not previously associated with somatic cell identity (grey) and prior markers of cell identity (light blue). (**m**) Heatmaps of the total (left), upregulated (middle left), and downregulated (middle right) number of pairwise differentially expressed genes. Right, example of gene expression Z-scores (estimated using all clusters) for differentially expressed genes between cluster 6 and cluster 7. (**n**) Number of differentially expressed genes (FDR < 0.05 and log_2_ fold-change > 0.25) for each pseudotime point versus T_0_ for each trajectory. (**o**) Left, log_2_ fold-change heatmaps between pseudotime points T_i_ and T_0_ for cluster 6 and cluster 7. Middle, heatmap of Pearson’s correlation coefficients (PCCs) between cluster 6 and cluster 7 log_2_ fold-changes across all pairwise timepoints. Right, average and 95% confidence interval of scaled PCC across all pairwise trajectories (top) and individual scaled PCC values derived from identical timepoints (bottom). The heatmap to right indicates the trajectories used in the comparison.

To better understand variation in transcriptional dynamics during dedifferentiation across trajectories, we modeled transcript abundance as a function of reverse pseudotime using generalized additive mixed models. In total, we identified 12,466 genes associated with dynamic expression across all dedifferentiation trajectories (FDR < 0.01; **Fig. S11b**). Of these, ∼24% (3,014 genes) were trajectory-specific, implicating heterogeneity in dedifferentiation regulatory programs (**Fig. 4d**). However, aggregating pseudotime-resolved gene expression patterns by the frequency of overlap revealed that genes most frequently shared across trajectories are biased for early onset expression, suggesting that a common set of genes initiate dedifferentiation (**Fig. 4d**). To test this, we performed pairwise global gene expression alignments between trajectories with distinct somatic and dedifferentiated cluster endpoints using dynamic time-warping (**Fig. 4e**). We found that the early and late stages of dedifferentiation were enriched for global alignments, consistent with a shared regulatory architecture independent of initial somatic cell identity and eventual dedifferentiated cell fates (**Fig. 4f**). To better clarify the factors underlying the early and late stages of dedifferentiation, we performed pairwise single gene alignments among all unrelated trajectories and defined aligned genes using empirical false discovery of permuted trajectories (eFDR < 0.05; **Fig. 4g**). Although the aligned trajectories were derived from different somatic origins, distinct dedifferentiated cell fates, and no overlapping cells (**Fig. S11c**), we found an average of 954 genes (range = 461-1,415) with statistically similar pseudotime-dependent patterns of transcription. Expression residuals for aligned genes partitioned by early, middle, and late stages of dedifferentiation were lowest in the early stages, consistent with a conserved dedifferentiation initiation program (Wilcoxon rank sum test, *P <* 2.2e^-16^; **Fig. S11d**-**S11e**).

Next, we aimed to characterize the generalized temporal hallmarks of dedifferentiation. To do so, we identified and *k*-means clustered all aligned genes (n=3,978 genes) based on averaged reverse pseudotime expression dynamics (**Fig. 4h**). Estimates of Shannon entropy indicated that temporal gene expression profiles were similar in all trajectories, despite most genes failing to align in more than 25% of trajectories (**Fig. S11f**-**S11h**). However, assessment of entropy scores across genes indicated an early period of reduced transcriptional heterogeneity followed by diversification of gene expression with progressive dedifferentiation (**Fig. S11f**). To define the shared processes at each stage of dedifferentiation, we performed GSEA across *k*-means clusters (**Fig. S11i**). GSEA exemplified the time-dependent signatures of protoplast dedifferentiation: (cluster 8) plastid and cellular reorganization; (cluster 7) ribosomal biogenesis and increased translation; (cluster 6) initiation of transcriptional and female gamete cell identity reprogramming; (cluster 5) chromatin remodeling; (cluster 4 and 3) hormone signaling; (cluster 2) transition to photorespiration; and (cluster 1) an increase in stress and wounding signatures (**Table S9**). Thus, several sequential cellular processes generalize dedifferentiation for diverse somatic origins and in vitro cell fates.

We hypothesized that the stereotypical dedifferentiation program is initiated by the same set of regulators regardless of the somatic cell identity. To investigate this, we generated trajectory-specific dynamic gene regulatory networks and assessed concordance of regulators (TFs, chromatin remodelers, signal transduction, and phosphoregulators) and their target genes. We found that regulatory genes activated at the early stages of dedifferentiation exhibited the greatest number of concordant targets across trajectories (**Fig. S12a**). Estimates of target gene overlap among distinct regulators within ordered activation timing bins (*n*=300) indicated prevalent regulatory redundancy at the onset of dedifferentiation (**Fig. 4i**). Next, we ranked genes by dedifferentiation initiation regulatory potential, estimated using activation timing, the number of concordant target genes, and consistent expression dynamics across trajectories (**Fig. S12a**). This analysis revealed a total of 405 genes with effects on early dedifferentiation (dedifferentiation initiation score > 0), including a 3-fold enrichment of genes with known roles in dedifferentiation (known regulator mean = 0.12 vs. unknown regulator = 0.04) such as *WIND1/2*, *HSFA1A*, *Rap2.6L*, *ZAT6,* and *MP* (Wilcoxon rank sum test, *P<*5.89e^-9^; **Fig. 4j**-**4k**). Investigation of functional annotations indicated that genes with greater dedifferentiation initiation potential (Z-score >1) were associated with terms such as hormone stimulus (*HB6*, *IAA18*, *ARF2*), stress response (*ERF4*, *WRKY48*, *MYB74*), and embryo development (*NF-YC4/10/11*, *NF-YA9*, *BT1*, *EIN3*) relative to all regulators (Hypergeometric test, FDR < 0.05). Moreover, we found that genes with strong regulatory potential were depleted of somatic cell type markers (Chi-square test, *P* < 1.1e^-3^). The few somatic marker genes with non-zero effects on early dedifferentiation were generally weaker than non-somatic markers, with the exception of *MP* which specifies vascular identity during both embryogenesis and post-embryonic development (**Fig. 4l**)^40^. These data implicate a core set of regulatory factors that initiate the generalized dedifferentiation program independent of somatic cell identity.

We then asked how initial somatic cell identity constrains dedifferentiated cell fates despite extensive sharing of regulatory factors during the early stages of dedifferentiation. Supporting diverse dedifferentiation outcomes, we found clusters predominantly composed of non-somatic cell types were highly distinct; pairwise comparisons indicated an average of 4,536 (range = 1,204-9,516) differentially expressed genes (DEGs) between dedifferentiated clusters (FDR < 0.05; **Fig. 4m**). However, dedifferentiated clusters were not as distinct as somatic cell types (average pairwise DEG = 6,773), consistent with dedifferentiation constraining transcriptional diversity (**Fig. 2j**, **Fig. S12b**). The simplest explanations for diversity in dedifferentiated cell fates are (i) heterogeneity in initial transcriptional reprogramming, (ii) partial redifferentiation from a common cell state, or (iii) incomplete repression of somatic identity regulators. To test these hypotheses, we first estimated transcriptional divergence from the initial somatic state as a function of progressive temporal dedifferentiation for each trajectory. We found that mesophyll-derived dedifferentiated clusters displayed the greatest somatic divergence in terms of the breadth and velocity of differentially expression (**Fig. 4n**, **Fig. S12c**). Peaks of differential expression velocity were often found at the onset of dedifferentiation, reflecting consistent early bursts of transcriptional reprogramming (**Fig. S12c**). However, early reprogramming events were frequently followed by periods of more modest yet increased expression changes (**Fig. S12c**). To establish the timing of in vitro cell fate diversification of mesophyll-derived cells, we correlated log_2_ fold changes vs. T_0_ at each subsequent time point between all pairwise mesophyll dedifferentiation trajectories. We found that early reprogramming changes are heterogeneous and exemplified by frequent, antagonistic, and small gene expression changes, despite all mesophyll dedifferentiation trajectories initiating from the same cell (**Fig. 4o, Fig. S12d**-**S12e**). In contrast, the later stages of dedifferentiation exhibited more consistent transcriptome-wide changes relative to the somatic mesophyll initial across trajectories (**Fig. 4o**). In parallel, somatic identity markers were strongly down-regulated in mesophyll but not guard cell or epithem-derived trajectories (Permutation, *P-*value < 1e^-4^; **Fig. S12e**). Thus, these data suggest that cellular diversity at the later stages of dedifferentiation is associated with small but opposing effects early during dedifferentiation, the timing of differential gene expression velocity peaks, and consistent loss of somatic marker expression.

## DISCUSSION

Understanding the regulatory mechanisms underlying dedifferentiation and the impact of somatic cell identity has remained a major question in developmental plant biology. Here, we leverage time course scRNA-seq of 125,091 Arabidopsis protoplasts derived from various seedling tissues and organs to understand transcriptional changes hallmarking dedifferentiation, which we define as the transcriptional deviation from somatic cell identities observed in Day 0 protoplasts. It is important to note that these data describe the early stages of dedifferentiation and are not predictive of cells that generate microcalli or have the capacity to regenerate. Although we may enrich for competent cells via strict data quality filtering, for example, by removing low quality barcodes, less than 1% of protoplasts ultimately gain regenerative competency. Instead, we asked how initial somatic cell states, hormone treatments, and culture time impact transcriptional reprogramming of in vitro cell suspensions, providing a much needed resource to better understand early dedifferentiation. We observed generalized transcriptional changes in cultured protoplasts consistent with prior observations, such as downregulation of canonical developmental programs, repression of photosynthesis, enhanced ribosomal biosynthesis and translation, and expression of TFs hallmarking dedifferentiation (e.g. *WIND1*) and stem cell identities (e.g. *LEC2*, *PLT7*, *ESR1,* etc.). We found that several markers of somatic embryogenesis and meristem identity are expressed heterogeneously immediately following protoplast isolation in a manner that depends on the initial somatic cell type. For example, ectopic expression of *LEC2* was restricted to cells with guard-like identity. Prior research has shown that epidermal cells in the stomatal lineage carry embryonic potential that can be realized by induced *LEC2* expression^37^. We speculate that the chromatin environment favorably positions these cells for somatic embryogenesis under in vitro conditions. More broadly, our results suggest that cell wall integrity signal transduction mechanisms are cell context dependent and constrain stochasticity in gene expression responses. Future studies of cell wall mechanosensing in diverse cell types would be informative for resolving the molecular cell-specific mechanisms underlying transcriptional responses necessary and sufficient to successfully reprogram totipotent cell identities.

In addition to the well-appreciated role in stimulating cell cycle re-entry, our analyses also suggest phytohormones contribute to secondary regulatory programs that promote dedifferentiation. By comparing protoplast cultures with and without auxin and cytokinin hormones, we found that phytohormone treatments antagonize cellular aging. Attenuation of cell aging coincided with cell state and transcriptional diversity, delayed signatures of senescence, and repression of JA wounding responses. Suppression of JA responses in hormone-treated cultures suggests a hormone feedback loop; JA spikes are known to promote auxin biosynthesis and de novo organogenesis in wounding experiments^41^. Analysis of dedifferentiation trajectories across distinct initial somatic cell types revealed that phytohormone applications strongly influenced early transcriptional dynamics and were associated with diversity in dedifferentiation outcomes. For example, application of phytohormones in somatic mesophyll cells promoted acquisition of distinct dedifferentiated states reminiscent of the known root reprogramming pwathway^23^, while non-hormone treated mesophyll cells expressed genes defining female embryo sac egg cell identity. These results suggest that exogenous phytohormone treatments, while not required for dedifferentiation, influence cell fate decisions. Long term culture experiments stratified by hormone treatments and dosages will be useful to understand the role of phytohormones in promoting disparate cell fate outcomes and the relationship with totipotency for a range of starting somatic cell identities.

Analysis of reverse pseudotime trajectories across initial somatic cell types indicated biases in dedifferentiation potential, the emergence of heterogenous dedifferentiated clusters, and a generalized regulatory program underlying dedifferentiation. Somatic cell type bias in dedifferentiation potential was consistent with prior studies of division-competent protoplasts in alginate embeddings, highlighting the accuracy of our probabilistic trajectory construction approach^15^. While mesophyll and guard-like cells have been previously implicated in protoplast regeneration^15^ and transgene-based somatic embryogenesis^37^, respectively, the dedifferentiation potential of epithem cells has not yet been described. Somatic epithem cells in this atlas uniquely express *YUC1,* which has been previously shown to induce MYB3R4-dependent cell cycle reactivation^15^. Our study indicated that withholding phytohormones in epithem cells promotes dedifferentiation, indicating that the total auxin concentration is an important determinant that shapes dedifferentiation potential. We posit that our experimental design of varied phytohormone treatments enabled the identification of epithem dedifferentiation missed by past studies, which have historically treated phytohormones as a fixed variable. These results suggest that distinct somatic cell identities may require tailored phytohormone concentrations for maximal dedifferentiation potential. Investigating the interplay between exogeneous and endogenous auxin and cytokinin concentrations and ratios across distinct somatic cells is an intriguing area of future study.

While our analyses have provided renewed insight into the regulatory processes underlying diversity in dedifferentiation, this data set also represents a valuable resource for future investigations. The atlas-scale profiling of diverse Arabidopsis tissues and organs can be leveraged for expedited cell type annotations in Arabidopsis seedlings across a range of genetic, chemical, and mechanical perturbations. The time-resolved nature of this resource provides a unique framework for anchoring single-cell data from additional developmental series and the ability to pinpoint instances of cell-specific dedifferentiation programs in canonical and perturbation contexts. Moving forward, we anticipate that expanding the temporal time course of these data along with tracking single cell transcriptomes with cell-specific phenotypes will be instructive for the rationale design of totipotency in plant cell culture applications. Maintaining dedifferentiated plant cells in culture is a critical first step towards establishing plant cell lines with controllable differentiation properties, enabling a wide array of stem cell-specific biological questions, and unlocking innovation in plant biotechnology.

## METHODS

### Plant growth conditions

*Arabidopsis thaliana* Col-0 seeds were obtained from the Arabidopsis Biological Resource Center and surface sterilized with a 10-minute incubation in a 50% bleach and 0.1% Sodium dodecyl sulfate solution, followed by five washes with sterile ddH_2_0. Surface sterilized seeds were sown on sterilized half-strength MS media (2.2g/L MS salts, 0.7g/L MES, 10g/L Sucrose, 5.5 g/L Agar, pH 5.6), stratified at 4 °C for 3 days, and grown under long-day conditions (16 hours light/8 hour dark at 22 °C) for 11 days.

### Protoplast isolation and culturing

Protoplast generation and culturing generally followed a pre-existing protocol previously demonstrated to induce plantlet regeneration^13^. Briefly, approximately 630 *Arabidopsis thaliana* Col-0 11-day old seedlings per biological replicate were submerged in 30 mL sterile 0.5M Mannitol solution (pH 5.8) and incubated at room temperature (RT) for 1 hour in a Petri dish. The Mannitol solution was then replaced with 30 mL of cell wall digestion enzyme solution (0.5% Cellulase R10, 0.4% Macerozyme R10, 0.1% Bovine Serum Albumin [BSA], 5mM 2-Morpholinoethanesulfonic acid, 5mM Calcium nitrate, and 500 mM Sucrose) filtered through a 0.22 μm syringe filter, followed by dark incubation at RT for 4 hours with gentle shaking at 15 rpm. Released protoplasts were passed through a sterile 40 μm cell strainer, washed by the addition of 30mL of washing solution (440 mM Mannitol and 0.1% BSA), filtered through a 40 μm cell strainer, distributed (∼15 mL) into sterile 30 mL round bottom tubes, and centrifuged at 300g for 5 minutes. After discarding the supernatant, 15 mL of washing solution was added to resuspend protoplasts with gentle shaking, followed by centrifugation at 300g for 5 minutes. Protoplast washing was repeated for a total of two washes. The supernatant was removed, and protoplasts were resuspended in 10 mL of washing solution. The preceding steps were performed in a sterile manner using 10 mL serological pipette tips.

Except for Day 0 samples, protoplasts at a final concentration of 1 x 10^5^ cells/mL were then dark incubated at RT in protoplast induction media (**Table S10**)^13^ with and without phytohormones (1 mg/mL 2,4-dichlorophenoxyacetic acid and 22ug/mL 6-Benzylaminopurine) for two, four, or six days prior to single cell RNA sequencing. Cell viability was estimated by treating a small aliquot of protoplasts with half-strength Trypan Blue stain.

### Culturing and imaging of reporter protoplasts

Seeds of Arabidopsis thaliana ecotype Col-0 expressing nuclear Red Fluorescent Protein (RFP) and membrane Yellow Fluorescent Protein (YFP) markers^42^ were sown on half-strength Murashige and Skoog medium containing 1% sucrose, stratified at 4 °C for 3 days, and grown under long-day conditions (16 hours light/8 hour dark at 22 °C) for 11 days. Whole seedlings were then harvested for protoplast isolation using the procedure described above with slight modification. Following addition of the cell wall digestion enzyme solution, samples were subjected to dark vacuum infiltration (30 minutes). After treatment, the samples were incubated for 4 hours at 23 °C with gentle shaking during incubation at 15 rpm. After the cell wall digestion, the enzymatic protoplast suspension was split and filtered through a 70 µm sterile mesh in a round-bottom tube, washed with 15 ml of pH 5.6 MES-based protoplast wash buffer (MES pH 5.7, mannitol 0.5 M, CaCl_2_ 10 mM, and 5 mM KCl) and centrifuged for 7 min at 100g, the supernatant removed, 15 ml of wash buffer added and the protoplasts resuspended by gentle shaking, centrifuged at 100g for 5 min; repeated for a total of three washes. After the final centrifugation, the protoplast suspension was diluted in protoplast induction media to a concentration of 1 x 10^5^ and cultured in the dark at 24 °C as described previously^16^. After 11 days, the medium was diluted to twice the volume with callus inducing media 1 (CIM1)^16^, again with (0 mg/mL 2,4-D and 0.11 mg/L 6-BA) and without hormones. After one month, the medium was further diluted to four times the original volume using CIM2 treatment medium, again with (0 mg/L 2,4-D and 0.22 mg/L 6-BAP) and without hormones. Two months later, the resulting callus (microcallus) was transferred to solid SIM medium containing hormones (0.1 mg/L IBA and 0.2 mg/L mT), with a 16 h light/8 h dark cycle at 24 °C. All media were adjusted to pH 5.6.

The excitation and detection setups for YFP and RFP were conducted as previously described ^43,44^. All optical micrographs were taken using a Zeiss LSM 880 Confocal Imaging System with Airyscan. For YFP, the emission was collected by a 543-615 nm filter. For RFP, the emission was collected by a 605.71-641 nm filter.

### Automated quantification of protoplast division

To quantify protoplast division, dual-channel z-stacks (RFP, nuclei; YFP, plasma membrane) were collapsed into channel-specific maximum-intensity projections. The final dataset comprised 131 fields distributed across 54 independently imaged areas (3–7 areas per day and condition; 2–6 fields per area). Cells were segmented using *Cellpose* v3 with the cyto3 model^45,46^. Segmentation was performed on the two-channel projections, with the membrane signal specified as the cytoplasmic channel and the nuclear signal as the nuclear channel. Cell diameter was estimated automatically, and relaxed segmentation thresholds (cellprob_threshold = −2.0; flow_threshold = 0.8) were used to improve recovery of cells with weak or incomplete membrane labeling. Raw 16-bit fluorescence intensities were provided without prior intensity transformation or normalization, such that only *Cellpose’s* internal image normalization was applied^45^. Nuclei were segmented independently by applying the same cyto3 model to the RFP projection alone ^46^. Candidate cell masks were retained if their area ranged from 50 to 25,000 pixels and their eccentricity was ≤0.93^47^. Individual nuclei were assigned to the cell mask containing the nuclear centroid^47^, and cells were subsequently classified according to nuclear number (0, 1, 2, or ≥3 nuclei). The division index was defined as the proportion of nucleated cells containing exactly two nuclei. Cells lacking a detected nucleus were excluded from the denominator. Multiple fields acquired from the same imaged area were pooled to calculate a single area-level division index, and imaged areas were treated as the unit of replication. Hormone-treated and untreated conditions were compared separately within each day using a two-sided Mann–Whitney U test^48^. Image handling, projection and plotting were performed in Python 3.12 using NumPy^49^, scikit-image^47^ and Matplotlib^50^.

### Construction and sequencing of single-cell RNA-seq libraries

Single-cell RNA-seq libraries were constructed following manufacturer instructions (10X Genomics) using the GEM single cell 3’ v3.1 (libraries 1-8) preparation kit (CG000315_ChromiumNextGEMSingleCell3_GeneExpression_v3.1_DualIndex RevE) and the GEM-X single cell 3’ v4 (libraries 9-19) preparation kit (CG000731_ChromiumGEM-X_SingleCell3_ReagentKitsv4_UserGuide_RevB) across three batches. For v3.1 kits, cell concentrations were adjusted to target recovery of between 3,000 and 10,000 cells (300-700 cells/μL). Cell concentrations (850-2,500 cells/μL) for v4 library preparations targeted recovery of 20,000 cells. scRNA-seq libraries were sequenced across two runs with 150bp paired end reads on an Illumina NovaSeq X.

### Initial scRNA-seq quality control

FASTQ files were processed and aligned to the TAIR10 Arabidopsis reference genome using 10X Genomics Cell Ranger (v9.0.1)^51^. Barcodes with less than 1,000 UMIs, 100 expressed genes, and 700 nuclear transcripts were removed from the analysis. Barcodes with fraction of chloroplast and mitochondrial reads greater than 2 standard deviations from the library mean and barcodes with fraction of transcript-derived UMIs less than 2 standard deviations from the library mean were removed. Although GEM-X technology can recover ∼20,000 cells at negligible doublet rates, only the top 16,000 barcodes ranked by total UMI counts were retained if more than 16,000 barcodes passed quality control filtering.

### Filtering scRNA-seq empty droplets and low-quality cells

Barcodes associated with low quality or background signal were removed by applying a previously described approach^52^. Briefly, we compared each step one filtered barcode (up to 16,000 per library, see above) to two aggregated references: (i) good quality cells defined as the top 10% of barcodes (minimum of 500 barcodes) ranked by UMI counts and (ii) background noise (i.e. empty droplets) defined as barcodes with less than 500 UMIs. For both references, the number of UMIs per gene were summed across all barcodes. Read counts across all genes were then scaled per 10,000 and log transformed for each reference. Highly variable genes from the good quality reference were identified by selecting genes with positive residual variance from a loess regression of per gene variance conditioned by the per gene mean across the top 10% (minimum of 500 barcodes) of barcodes. We then estimated Pearson’s correlation between individual barcode normalized expression values (scaled per 10,000 and log transformed) and the normalized expression of the good quality and background noise references conditioned on the highly variable gene set. Barcodes with a higher correlation to the good quality reference compared to the background noise reference and raw Pearson correlation coefficients to the good quality reference greater than 0.075 were retained.

### Identification of scRNA-seq multiplet barcodes

Barcodes passing the above quality control and background filtering steps above were screened for potential doublets using *DoubletFinder* v2.0.4^53^. Briefly, the barcode meta data and UMI counts matrix for each library was loaded into a *Seurat* (v5.0.1)^54^ object and processed with default normalization and dimensionality reduction settings (*SCTransform*, *RunPCA*, *RunUMAP*, *FindNeighbors*, *FindClusters*). The optimal DoubletFinder parameter for neighborhood size (pK) was identified as the maxima of the mean-variance normalized bimodality coefficient using the functions, *paramSweep* (sct=TRUE), *summarizeSweep* (GT=FALSE), *find.pK* with non-default settings. The expected proportion of homotypic doublets was determined by adjusting the expected doublet rate based on loading concentration (conservatively set at ∼10%) with one minus the output of *modelHomotypic* with default settings. Doublet barcodes were then identified using *doubletFinder* with non-default settings (obj, PCs=1:50, pN=0.25, pK=best.pk, nExp=nExp_pois.adj, sct=TRUE). Only barcodes classified as singlet cells were retained for downstream analyses.

### Single-cell RNA-seq clustering

To characterize plant cell dedifferentiation, we first constructed a cell x gene counts matrix using the Day 0 protoplasts, which should largely exhibit differentiated cell types. After removing chloroplast and mitochondrial genes, we identified and excluded outlier cells (top 99% quantile) based on total UMI and expressed gene counts. Counts normalization and clustering was performed with the R package, *Seurat* (v5.0.1)^54^. Briefly, the counts matrix was log transformed using the default settings of *NormalizeData,* variable genes identified with the default settings of *FindVariableFeatures,* and centered expression values generated by the default settings of *ScaleData.* The top 50 principal components (PCs) were identified with the function *RunPCA* with non-default settings (approx=F, npcs=50). Batch effects were removed with the R package, *harmony* v1.2.0^55^ with non-default settings for the function, *RunHarmony* (obj, “library”, theta=2, nclust=100, max.iter=30). The corrected PCs were used as input for the function *umap* from the uwot R package with non-default settings (min_dist=0.3, n_neighbor=30, metric=”correlation”). A shared nearest neighbor graph was constructed using the function, *FindNeighbors* with non-default settings (reduction=”harmony”, k.param=30, dims=1:30, prun.SNN=1/30, l2.norm=T). Initial clusters were identified with *FindCluster* with non-default settings (resolution=0.8, algorithm=1). Four of the major clusters exhibited mixed marker gene expression profiles. For these clusters, we performed an additional round of sub-clustering with the function *FindSubCluster* with non-default settings, where we selected the resolution for the focal cluster that best partitioned mixed marker expression patterns (resolution=res[i], cluster=cl[i], algorithm=4).

Data processing and clustering for the full data set (Day 0, 2, 4, and 6) was performed similarly as the Day 0 protoplast samples. Briefly, we removed chloroplast and mitochondrial genes, and outlier cells (top 99% quantile) based on expressed genes and total UMI counts. The counts matrix was log normalized and centered using *NormalizeData* and *ScaleData,* respectively, with default settings. The top 50 PCs were identified using *RunPCA* (approx=F, npcs=50). Batch effects were removed with *RunHarmony* with non-default settings (obj, c(“replicate”, “batch”, “tech”), theta=c(1,1,1), lambda=NULL, nclust=100, max_iter=30). Corrected PCs were projected in the UMAP space with *umap* with non-default settings (min_dist=0.3, n_neighbors=30, metric=”correlation”). The shared nearest neighbors graph was constructed using the function, *FindNeighbors* with non-default settings (reduction=”harmony”, k.param=30, dims=1:30, prun.SNN=1/30, l2.norm=T). Finally, initial clustering was performed with *FindClusters* with non-default settings (resolution=1.5, algorithm=1).

### Single-cell RNA-seq annotation

To annotate differentiated cell types from Day 0 samples, root and leaf cell type marker genes were first collected from references^29,56^. Next, we estimated cluster specificity for each gene using the tau metric^57^. All cell type marker genes from scPlantDB were ranked by significance and log_2_ fold-change within each individual data set, respectively. We then estimated a combined specificity score for each cell type marker gene using equation (eq. 1):

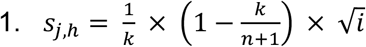

Where *s_jh_* is the combined specificity score of gene *j* for cell type *h*, *k* is the number of data sets where marker *j* was significant (FDR < 0.05) for cell type *h*, *n* is the total number of independent data sets that include cell type *h,* and *i* is the average rank of marker gene *j* in cell type *h* across all independent data sets. The top 5 ranked markers based on the combined specificity score, FDR < 0.05 across multiple independent data sets, positive (PCC > 0) co-expression correlation with other marker genes for the same cell type, and tau cluster specificity metrics above 0.6 were retained. Gene expression signatures for the curated set of marker genes were visualized on UMAP embeddings after smoothing log normalized expression values via the MAGIC algorithm (10 nearest neighbors and 5 steps)^58^, as well as depicted as standardized values (Z-scores) using cluster-averaged log normalized expression. Day 0 cell type annotations were determined manually by reviewing marker expression patterns from both visualization techniques.

### Differential gene expression among clusters

Differential gene expression analysis among clusters from the Day 0 only and integrated analyses were conducted similarly. Briefly, the log normalized counts were used as input for the *Seurat* function, *FindAllMarkers* with non-default settings (logfc.threshold = 0.25). Genome-wide significance was determined using Bonferroni-correct *P-*values less than 0.05 and an absolute log_2_ fold-change greater than 1.

### GO term enrichment

Up- and down-regulated differentially expressed genes from the Day 0 cell-depleted clusters were uploaded individually to the GO enrichment tool provided by the online PANTHER database (v19.0). The reference list was set to the union of all expressed genes across all protoplast culture duration and phytohormone-treated samples. *P-*values were calculated using Fisher’s exact test and multiple-test corrected using Benjamini-Hochberg FDR. The complete GO biological process list was used as the annotation data set. GO terms with FDR < 0.05 were selected as significantly enriched or depleted.

### Differentiation scores

Per cell differentiation scores were calculated by comparing each cell from the integrated embedding with Day 0 pseudocells generated by this study or with somatic cell type clusters defined by Tenorio Berrio et al^29^. Briefly, log normalized expression values from each cluster were averaged across member cells. We then estimated Pearson’s correlation coefficient between the log normalized expression values for each individual cell and the somatic pseudocells using the function, *corSparse* from the R package, *qlcMatrix*. Given a cell by somatic pseudocell correlation matrix, differentiation scores were then defined as the maximum Pearson’s correlation coefficient value across all differentiated pseudocells for a given cell. Day 2, 4, and 6 samples with differentiation scores greater than the 5% quantile of Day 0 differentiation scores (Pearson’s correlation coefficient of ∼0.5) were annotated with the corresponding cell type and “dedifferentiated” otherwise.

### Gene expression patterns associated with differentiation

We implemented two complementary approaches for associating patterns of gene expression with differentiation scores. First, we calculated Pearson’s correlation coefficients between the log normalized expression levels and differentiation scores for each gene. To determine significance, we scrambled cell IDs of the cell by gene expression matrix, and recalculated Pearson’s correlation coefficients. Permuted PCC scores were conducted 1,000 times to generate a null distribution for each gene. PCC-based Z-score were then estimated by mean subtraction and division by the standard deviation of the null distribution with respect to the observed PCC estimate from the original matrix.

Linear mixed effects models (LMMs) have been shown to effectively control false positive associations stemming from non-independence among cells of an individual. Using this approach, we modeled per cell differentiation scores as a function of library ID coded as a random effect, and technical metrics including batch, log UMI, and log normalized expression level as fixed effects for each gene with non-zero counts in at least 50 cells. The full model was compared to a reduced model lacking a coefficient for gene expression using Analysis of Variance (ANOVA). Specifically, we estimated and compared the differentiation score fits of eq. 2 and eq. 3 for each gene assuming a Gaussian error distribution.

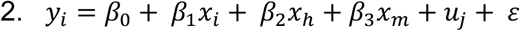

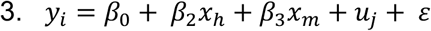

Where *y_i_* is a vector of differentiation scores across cells, *β*_0_ is the intercept, *x_i_* is a vector of log normalized expression levels of gene *i*, *x*_ℎ_ is a vector specifying the log UMI sum of each cell, *x_m_* is a vector containing batch assignment of each cell, *u*_j_ is the random effect of library *j,* and *ε* is the residual error. *P-*values were derived from the Chi-square distribution by comparing the fits of eq. 2 and eq. 3 with a likelihood ratio test for each gene by way of ANOVA. Adjusted *P-*values were estimated using Bonferroni multiple test correction with the R function, *p.adjust* with non-default settings (method=”bonferroni”). Genes with adjusted *P-*values less than 0.05 were identified as significantly associated with differentiation.

### Modeling cell age in real time from cellular state

To assign precise cell ages in hormone and non-hormone treated samples, we trained regularized elastic net models to learn sample age from individual cell transcriptomes, split by hormone treatment status. Specifically, the log-normalized expression counts matrix was partitioned into two groups, (1) hormone-treated cells, and (2) non-hormone treated cells. Since day 0 cells were used as the source of both hormone and non-hormone cultures, the transcriptomes of day 0 cells (4 hours old protoplasts) were included in both the hormone-treated and non-hormone treated digital gene expression matrices. Thus, both hormone and non-hormone models were trained with the same day 0 cells, while day 2 and day 4 cells were unique to the hormone treatment. Day 6 cells from the hormone-treated gene expression matrix were removed to include only matched protoplast ages in both hormone and non-hormone groups. To account for class imbalance, the number of cells from each age group were down sampled to match the protoplast age with the fewest number of cells across both treatment types (*n* = 5,560 per protoplast age group in both treatment types). Training and test cell identifiers were identified using the function *createDataPartition* from the R package, *caret* v7.0.1 with non-default settings (p=0.75, list=F, times=1)^59^. Elastic net models were trained on the transcriptomes of hormone and non-hormone cells with the function *cv.glmnet* from the R package, *glmnet* v4.1.8 with non-default settings (alpha=0.5). The models were used to predict cell age in the 25% of held out cells, as well as the full data set, using the *predict* function with non-default settings (s=’lambda.min’, type=’response’). Predicted age accuracy was determined by Pearson’s correlation coefficient between predicted and ground truth values.

### Gene set enrichment across real time

To identify changes in aggregate marker gene expression across time, we collected genes associated with aging (GO:0007568) and senescence (GO:0010149) gene ontology terms. To determine per cell enrichment scores by gene set, we first calculated the fraction of marker genes expressed in the cell. We then built null distributions matched to the gene set of interest by randomly selecting the same number of genes randomly and estimating the fraction expressed genes in the random set across 100 permutations. Gene set enrichment scores per cell were then estimate by standardizing the observed fraction of expressed marker genes (*x*) using the mean (*u*) and standard deviation (σ) from the null distribution (eq. 4).

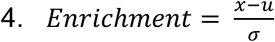

We then used the gene set enrichment scores as the response variable in generalized additive models fit with smoothed representations of real time predictions as the explanatory variable using the function *gam* from the R package, *mgcv*^60^ with non-default settings (enrichment ∼ s(real_time, bs=”cr”)). The mean fit and standard deviation was estimated via the *predict* function (model, newx, type=”link”, se.fit=T).

### Phytohormone-associated differential cell state abundance and gene expression across real time

To make use of the full data set for testing differential cell state abundances and gene expression between hormone and non-hormone treatments, the same model architecture and prediction validations as above were used with the exception of including day 6 cells in hormone-treated model training. All other parameters, such as 3:1 training to testing partition, 10-fold cross-validation, class balancing, and prediction arguments were identical.

For testing differential cell state abundance, cell meta data, log-normalized and library size-adjusted gene expression values, and all reduced dimension embeddings were imported into a SingleCellExperiment object compatible with the R package, *SingleCellExperiment* (v1.20.1)^61^, and then converted into a *Milo* object using the function *Milo* from the R package, *miloR* v1.6.0, with default settings^62^. Graph construction and neighborhood assignment was performed with non-default parameters with the functions *buildGraph* (k=20, d=20, reduced.dim=”HARMONY”) and *makeNhoods* (k=20, d=20, refined=T, prop=0.2), respectively. Neighborhood connectivity and cell counts within neighborhoods across replicated conditions was conducted using non-default parameters of the functions *calcNhoodDistance* (d=20) and *countCells* (samples=”library”, meta.data=colData(sce)), respectively. The experimental design matrix for differential abundance testing was constructed using hormone treatment as the explanatory variable and library identifier as a covariate with the function *xtab* (∼ hormone + library, data=colData(sce)). Identification of differential cell state abundance associated with hormone treatment was conducted using the function *testNhoods* with non-default parameters (design= ∼hormone, design.df=design.matrix). Significant differences in cell state abundances due to hormone treatment for each neighborhood were determined using a spatial FDR threshold of 0.05.

Dynamic gene expression conditioned by hormone treatment was modeled using two generalized additive mixed models (GAMMs) fit to each gene, *i*. Briefly, two GAMMs were fitted to assess how gene expression (*y*) varied with technical and biological covariates. Both models were implemented using the *mgcv* package in R, with smoothing terms estimated via restricted maximum likelihood (REML) to balance model fit and smoothness. The first model (eq. 5) included fixed effects for sequencing technology (tech), batch (batch), total UMI counts (log_umi), and hormone treatment (hormone). To capture potential nonlinear relationships between predicted cell age and gene expression that differ by hormone treatment, the model incorporated a smooth functional term of predicted cell age stratified by hormone treatment [s(inferred_age, by = hormone)]. A random effect smooth on library [s(library, bs = “re”)] was included to account for non-independence among cells derived from the same biological sample. Formally, let *y_i_* be a vector of log normalized expression values for cell *i*.

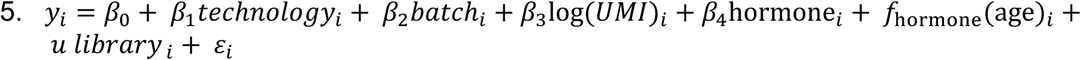

Where *f*_hormone_(age)*_i_* is a hormone-specific smooth function of predicted cell age modeled using a penalized spline and *u library _i_* is a random intercept to account for non-independence among cells from the same biological sample. We also fit a second nested model (eq. 6) lacking a fixed effect for hormone treatment and modeled predicted cell age independent of hormone treatment.

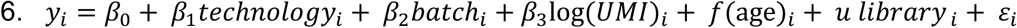

After extracting model coefficients, each model was refit using maximum likelihood. We then compared the full and nested models using a generalized likelihood ratio test (ANOVA with a Chi-square test method) to estimate a *P-*value for the effect of hormone on the smoothed gene expression patterns. *P*-values were adjusted for multiple testing using *p.adjust* with non-default settings (method=”fdr”). Genes with FDR < 0.05 were considered significant. To improve computational efficiency, this approach was implemented on a reduced data set where each library was first down sampled to a maximum of 1,000 cells.

### Integration and comparison of snRNA-seq and scRNA-seq

To evaluate transcriptome changes unique to protoplasting, we integrated our 11-day Arabidopsis seedling protoplast-based scRNA-seq data with previously published snRNA-seq data derived from 12-day Arabidopsis seedlings^63^ using the R package, *rliger* (v2.2.1)^64^. Briefly, both datasets were merged into a Liger object using the function *createLiger* with non-default parameters (list(nuclei=sn, protoplast=sc)). Cells with at least 700 UMIs and 300 expressed genes across both data sets were retained. Normalization, variable genes, and scaling (without centering) were performed with default parameters using the function *normalize, selectGenes, and scaleNotCenter*, sequentially. The two modalities were integrated using the functions *runIntegration* (k=30) and *alignFactors* (method=quantileNorm). Clusters were identified using the Leiden algorithm via the function *runCluster* with a resolution = 0.3. Differential expression between protoplast and nuclei cells within a cluster was performed using a pseudobulk differential expression framework via edgeR^65^. To do so, UMI counts for protoplasts or nuclei within a cluster from the same replicate were aggregated and filtered to remove pseudobulks with fewer than 100 cells or nuclei. Clusters with fewer than two replicates from both protoplast and nuclei-based were removed. Differentially expressed genes between nuclei and protoplast modalities were then identified using the standard edgeR workflow with default settings.

### Reverse pseudotime trajectories

To generate independent estimates of differentiation, the top 30 cell embeddings from the harmony integration were input to the function *DiffusionMap* from the R package, *destiny* (v3.21), with non-default parameters (n_eigs=30)^66^. However, individual diffusion components failed to accurately represent all possible somatic dedifferentiation trajectories and instead reflected global dedifferentiation, similar to the correlation-based differentiation score, necessitating an alternate approach. To this end, we applied *Palantir* (v1.4.1) to explicitly model trajectories of dedifferentiation for each somatic cell type in probabilistic framework^67^. We first generated diffusion maps using the integrated harmony cell embeddings using the function *run_diffusion_maps* with non-default parameters (knn=15, n_components=100). Start cells were identified by selecting the cell closest to the cluster centroid within the harmony embedding conditioned by the dominant treatment sample within the cluster. For example, we required that the cluster 4 start cell was selected from a Day 4 non-hormone treated library and cluster 6 and 18 start cells from the Day 6 hormone treated library. Start cells were selected from all clusters with more than 50% of cells classified as dedifferentiated, collectively representing 10 different stem cell clusters. Terminal cells were identified for each somatic cell type as the cell closest to the centroid of the cognate cell type within the integrate harmony embedding using all cells from the Day 0 libraries. Reverse pseudotime across the entire data set for each putative stem cell population (10 start cells x 21 stomatic cell types = 210 reverse trajectories) was estimated from the multi-scale low dimensional embedding based on the top diffusion components defined by an eigen gap. Reverse pseudotime estimates were calculated independently for each putative stem cell cluster start cell using the same terminal cells with the *run_palantir* function and non-default parameters (num_waypoints=2500, use_early_cell_as_start=True, terminal_states=terminal_states.index, n_jobs=1, knn=30). Consensus pseudotime was calculated as the minimum reverse pseudotime value across the 10 independent reverse trajectories for each cell.

### CytoTRACE-based estimates of differentiation potential

Stem cells generally exhibit greater transcriptional diversity compared to terminally differentiated and specialized cell types^35^. Thus, estimates of transcriptional diversity offers a complementary approach to evaluate diffusion-based pseudotime methods. To achieve this, the log-normalized counts matrix was filtered to remove genes expressed (log-normalized counts greater than 0) in fewer than 100 cells. The filtered log-normalized matrix was then used as input for the R package, *CytoTRACE* (v0.3.3) with non-default settings (batch=obj@meta.data$libraryID, subsamplesize=5000, enableFast=T).

### Guard cell time-lagged gene regulatory network construction

To identify putative regulators and targets of LEC2 in dedifferentiating guard cells, cells belonging to cluster 18, the cluster containing the majority of Day 0 cells with guard cell identity, were isolated from the log-normalized gene expression by cell identifier matrix. Genes with zero log-normalized expression across cluster 18 cells were removed. To limit analysis to genes putatively associated with time-resolved or hormone-dependent expression patterns, we modeled gene expression as a function of the interaction between real-time cell age and hormone treatment. To air on the side of more inclusive feature selection, genes with nominal *P*-values greater than 0.05 for all coefficients (age, hormone status, or age x hormone status) were removed. The filtered log-normalized gene expression by cell matrix was then ordered by diffusion pseudotime, ranking cells from somatic to dedifferentiated. Gene expression values were smoothed by average log-normalized expression in sliding windows of 7 cells with a one cell step separately for hormone and non-hormone treated cells. A first-order lag model (lag = 1) was used to infer dynamic TF-target gene associations. Specifically, for each cell *i,* TF expression at time *t-1* was used to predict target gene expression at time *t.* In this way, time-lagged matrices of TF regulator expression values were constructed for hormone and non-hormone treated cells independently. The first diffusion pseudotime cell in the time-lagged TF expression matrices for each condition was excluded as a result of unmeasured lag values.

Two types of dynamic gene regulatory networks (dGRNs) were generated from these input: (1) Condition-specific dGRNs, and (2) a dGRN testing interactions between TF expression and hormone treatments. In the first type of dGRNs, hormone and non-hormone treated cells were modeled independently using regularized ridge regression from the R package, *glmnet* (v4.1.8). Specifically, we fit a penalized linear model for each target gene *g* with eq. 7:

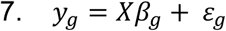

Where *y*_g_ is the smoothed target gene expression across diffusion pseudotime and *X* is the time-lagged TF gene expression matrix ordered by diffusion pseudotime. The models were fit with L2 regularization (ridge regression) by setting the alpha parameter in the call to *glmnet* equal to 0. The penalty parameter was selected using 5-fold cross-validation with internal standardization. To identify significant time-resolved associations between time-lagged TF gene expression and target genes, we adopted a permutation-based approach. A null distribution of target-regulator TF coefficients was generated by permuting the temporal alignment between predictors and response variables prior to ridge regression coefficient estimation for a total of 100 permutations per target gene. A z-score was then estimated by subtracting the mean of the null coefficient distribution from the observed coefficient and dividing by the standard deviation of the null distribution. Z-scores were then converted to *P-*values assuming a normal approximate. We also estimated empirical *P-*values using eq. 8:

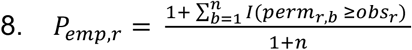

Where *perm_r_*_,*b*_ is the permuted coefficient of permutation *b* and *obs_r_* is the observed coefficient. Following *P-*value estimation for all TF regulator-target pairs, both the normal approximated and empirical *P-*values were corrected for multiple testing using the R function, *p.adjust* with non-default settings (method=”fdr”). TF regulator-target pairs with FDR < 0.01 were considered statistically significant.

To test for hormone treatment effects, the condition-specific TF time-lagged matrices were joined to generate a design matrix that included the lagged regulator smooth log-normalized expression values, hormone interactions terms, and the diffusion pseudotime covariate. Therefore, each target gene *i* was modeled using regularized ridge regression as shown in eq. 9:

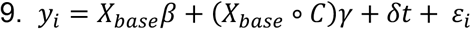

Where *X_base_* is the time-lagged TF regulator matrix, *C* is the binary condition indicator, *X_base_* ∘ *C* is the element-wise interaction term, *t* is the diffusion pseudotime vector, *β* is the baseline regulatory effects, and *γ* is the condition-specific deviations. Regularization of coefficients was performed by setting the alpha parameter in *glmnet* to 0 with 5-fold cross-validation to minimize the penalty parameter, *λ*, error. To identify significant interactions between hormone treatment and TF regulator-target associations, we implemented a permutation approach similar to the single-condition dGRNs. Rather than randomizing the temporal alignment, we generated the null distribution of interaction coefficients by permuting hormone and non-hormone labels among cells and re-estimating interaction coefficients across 100 permutations. Normal approximated *P-*values, empirical *P-*values, and cognate FDRs were calculated identically as in the single-condition dGRNs. TF regulator-target gene pairs with FDR < 0.01 were considered statistically significant.

### ATAC-seq, bulk RNA-seq, and TF motif analysis

Three replicates of bulk ATAC-seq and RNA-seq data corresponding to intact leaves and protoplasts (8 hour overnight release) generated with the same protocol were collected from a previously published study available on NCBI BioProject, PRJNA648028^16^. Raw ATAC-seq sequencing reads were trimmed using default settings of *fastp* (v0.23.4)^68^ and aligned to the TAIR11 Arabidopsis reference genome using the default settings of *bwa mem* (v0.7.17)^69^. PCR duplicates, non-properly paired, multi-mapped, and low mapping quality reads (MQ < 30) were removed. ACRs were identified using *MACS2* (v2.2.4) with non-default settings (--nomodel --keep-dup all --extsize 150 --shift -75 --qvalue 0.05 --bdg)^70^. Bedgraph coverages output by *MACS2* were converted to bigwig format using the *bedGraphToBigWig* tools from UCSC utilities (v4.0.0). To reduce bias in ACR identification due to technical variation (for example, in the analysis of ACR size changes), we also called ACRs in protoplast and intact leaves after concatenating cleaned reads across replicates and downsampling to a fixed number (80 million) reads for each sample. ACRs called (FDR < 0.05) from downsampled reads were used to assess ACR size differences and to define ACR sequences unique to protoplast conditions, intact leaves, and shared between protoplast and intact leaves. Differential ACRs were identified using *edgeR* (v3.40.2) using the union of all ACRs across the three replicates of intact leaves and protoplast ATAC-seq data^65^. Differential ACRs were defined as ACRs with FDR < 0.05. TF motifs within ACRs were identified using *motifmatchR* (GitHub version d0dc70e) with a *P-*value threshold of *P*-value < 1e^-5^. Differential motif enrichment was estimated using Fisher’s exact test between motif counts in protoplast-specific and shared ACRs, and intact leaf-specific and shared ACRs. Nominal *P-*values from differential motif enrichment were multiple test corrected using *p.adjust* with non-default settings (method=”fdr”). Condition-enriched TF motifs were defined as FDR less than 0.05 and a log_2_ fold-change relative to stable ACRs greater than 0. *TFBSTools* (v1.48.0) was used to visualize motifs from the noted WRKY TFs ^71^. Normalized bulk RNA-seq counts data between day 0 protoplasts and intact leaves was directly acquired from the previously published study without further processing^16^.

### Multiscale footprinting analysis

To identify TF and nucleosome occupancy changes between leaf and protoplast, we developed a library-equalized bulk footprinting approach. Intact leaf and protoplast BAM replicates (MAPQ ≥ 10, PCR duplicates removed) were subsampled to equal sequencing depth at the replicate level using *samtools view -bs* (seed = 42)^72^. Replicates were then merged per condition using *samtools merge* to produce two pooled BAMs with identical library sizes. Merged BAMs were name-sorted and converted to scPrinter-compatible 1-based fragment files using a custom extraction script (bulk mode, minimum MAPQ = 20), with 0-based to 1-based coordinate conversion. Mitochondrial and chloroplast reads were excluded. ACR classification used the edgeR results described above (FDR < 0.05), yielding 1,222 protoplast-enriched, 2,873 leaf-enriched, and 18,487 stable ACRs. All ACRs were center-extended to a uniform 2,000 bp width for batch processing, with one ACR dropped due to chromosome boundary clipping, yielding 22,581 resized regions. A native-to-resized coordinate mapping table was generated for downstream motif-to-region analysis.

To reduce within-family motif redundancy while preserving between-family diversity, we clustered 465 *Arabidopsis*-specific motifs from the JASPAR 2024 CORE plants collection into non-redundant signatures using the R package, *motifStack*^73^. Motifs were clustered independently within each TF family to prevent biologically distinct motifs from different families from merging based on superficial PWM similarity. Singleton families were passed through directly while pairs were merged if their clustering distance was < 0.5. Families with ≥ 3 motifs were clustered with non-default parameters via *motifSignature()* with *cutoffPval = 0.0001* and *min.freq = 1*. This produced 122 non-redundant signatures across approximately 42 TF families. Signatures were scanned on native (non-resized) ACR sequences using MOODS^74^ with a per-motif *P*-value threshold of 5 x 10^-5^, scanning both strands with per-motif *Scanner* instances. Position frequency matrices were converted to position weight matrices via log-odds transformation against a uniform nucleotide background (0.25) with a pseudocount of 1 x 10^-4^. The scan yielded 3,785,216 hits across 22,581 ACRs.

For each condition, *scPrinter* (v1.2.1)^39^ was applied to the 22,581 resized ACRs with standard Tn5 strand offsets (+4/-5 bp) and no coverage filters (*min_num_fragments = 0, min_tsse = 0*). Three models were computed per condition: (i) multiscale footprint scores (FP) at 99 scales (2–100 bp) via *get_footprint_score()* with *modes = np.arange(2, 101)* and *region_width = None*, representing −log_10_(*P*-value) significance at each scale and genomic position, producing a tensor of shape (1, 99, 2000) per region; (ii) TF binding score (TFBS) using the pretrained neural network model (*pretrained_TFBS_model*), which consumes footprint signals at 6 scales (10, 20, 30, 50, 80, 100 bp) within a ±100 bp context window, producing a sigmoid-activated probability in [0, 1] per tile; and (iii) nucleosome binding score (NucBS) using the pretrained nucleosome model (*pretrained_NucBS_model*), which uses 5 scales (10, 20, 30, 50, 80 bp) within the same context window. The NucBS model lacks an output activation function, producing unbounded raw scores; a numerically stable sigmoid transformation was applied post-hoc, using a two-branch formulation: σ(x) = 1/(1+e⁻ˣ) for x ≥ 0, and σ(x) = eˣ/(1+eˣ) for x < 0, to avoid floating-point overflow on extreme raw scores (observed range −10 to +6), thus converting raw scores to [0, 1] probabilities. Both binding score models use a tiling scheme of 180 tiles per region (*tileSize = 10, contextRadius = 100, downsample = 5*), with tile centers at positions 105, 115, …, 1895 bp within each 2,000 bp window. Both the TFBS and NucBS models were adapted from the original multiscale footprinting publication^39^. Because the models detect footprint shape patterns – the Tn5 insertion protection dip created by a bound protein has similar geometry across species – the models are presumed to be organism-agnostic. However, absolute probability values are not calibrated for *Arabidopsis* and were therefore treated as relative rankings rather than calibrated binding probabilities. In contrast, the multiscale FP score is a statistical test (−log_10_ *P*-value) using an *Arabidopsis*-specific Tn5 bias model and is fully valid for this organism.

To assess systematic condition bias across FP scales, we compared leaf−protoplast deltas at 22,338 ACR centers (position 1000 within the 2,000 bp window) and at 4,937 motif-free null loci. Null loci were sampled by selecting random positions within ACRs at least 50 bp from any JASPAR motif hit and at least 60 bp from ACR edges (seed = 0, target = 5,000). One-sample t-tests against delta = 0 were performed per scale with Bonferroni correction (α = 5.1 x 10^-4^ for 99 tests). No TF scale (2–20 bp) reached significance at null loci, while 27 of 99 scales at >40 bp showed modest but significant protoplast favoring bias (maximum |Δ| = 0.047), consistent with global chromatin reorganization rather than TF-specific signal. The full null delta distribution per scale was retained for downstream empirical Z-score correction.

To identify optimal diagnostic scales, per-scale Spearman correlations between binding score probabilities and FP signal were computed for tiles classified as bound/unbound (TFBS) or occupied/free (NucBS) using per-condition percentile thresholds. TFBS probability correlated most strongly with FP at 4–10 bp (TF protection range), while NucBS occupancy peaked at 40–60 bp (nucleosomal range). These correlations were consistent across conditions and across three threshold stringencies (top/bottom 2%, 5%, and 10%), confirming the adequacy of the corresponding scales to capture distinct, scale-appropriate chromatin features in both conditions. Tiles were then classified as bound (TFBS, top 2%, 5%, and 10%) or occupied (NucBS, top 2%, 5%, and 10%) using per-condition percentile thresholds computed across all regions and tiles. The percentile thresholds were chosen rather than absolute cutoffs because the score distributions are condition- and model-dependent. Percentile-based thresholds ensure equal representation of extreme tiles across conditions. Overlap between leaf and protoplast FPs was quantified per ACR class using hypergeometric tests on the universe of active tiles (bound or occupied in at least one condition).

For each of nine categories (3 ACR classes × 3 overlap groups: leaf-only, shared, protoplast-only), FP deltas (leaf minus protoplast) were extracted at TF scale (4–10 bp, 7 scales averaged) and nucleosome scale (40–60 bp, 21 scales averaged). Deltas were tested against a genome-wide null distribution derived from tiles classified as unbound/free in both conditions (bottom-percentile tiles in both leaf and protoplast), subsampled to a maximum of 50,000 tiles. Category-level significance was assessed by Mann-Whitney U test with Benjamini-Hochberg FDR correction across the 9 tests. Individual tiles were then Z-scored against the null: z = (Δ_tile – Null mean) / Null standard deviation. Tiles with Z-score ≥ 1 were classified as leaf-enriched, tiles with Z-score ≤ −1 as protoplast-enriched, with BH-FDR correction applied within each category. TF motif hits from the non-redundant signature database were mapped to tile coordinates by translating native ACR motif hit centers to resized coordinates and assigning each to the overlapping tile (tile center ± 5 bp). Hits were deduplicated by region × tile × family to avoid double-counting. For each of 18 combinations (9 categories × 2 directions: leaf-enriched Z-score ≥ 1, protoplast-enriched Z-score ≤ −1), Fisher’s exact test (two-sided) assessed whether specific TF families were enriched among tiles with significant FP deltas relative to the genome-wide background of all active tiles. The 2 × 2 contingency table compared foreground tiles (significant delta in the specified direction) versus background tiles (all active tiles genome-wide), crossed with presence or absence of a motif hit from the focal family. BH-FDR correction was applied per combination across all tested families.

### Refining dedifferentiation trajectories with probabilistic walks

To define precise paths between somatic cell states to stem cell like identities, we first constructed pseudotime-constrained probabilistic graphs of cell state transitions using a *k* nearest neighbors approach. Dedifferentiated cells were first partitioned into three groups based on their maximum cell origin probability (spongy mesophyll, guard cell, or epithem cell) and concatenated with Day 0 cells of the same cell type. To reduce the search space to only likely paths between somatic cell types and the 10 stem cell clusters, we required that a path between a somatic cell type and a dedifferentiated cluster have at least 100 cells in the dedifferentiated cluster contain a maximum cell origin matching the somatic cell state (i.e. at least 100 dedifferentiated cells in cluster 1 with maximum cell origin of spongy mesophyll for assessing a path from somatic mesophyll cells). This criteria resulted in a total of 12 possible dedifferentiation trajectories: 10 spongy mesophyll trajectories, one guard cell trajectory, and one epithem trajectory. For each of the 12 possible dedifferentiation trajectories, we then identified the 30 nearest neighbors among all cells with the same maximum cell origin probability or day 0 somatic cell type label using the top 100 diffusion components output by *Palantir* (i.e. day 0 spongy mesophyll cells and day 2+ dedifferentiated cells with maximum cell origin labels equal to spongy mesophyll). Starting from the same initial cell coordinate as for reverse pseudotime estimation in *Palantir,* each knn graph was then iteratively refined to only keep edges between cells with equal or increasing pseudotime values. This results in a directed graph constrained by progressive pseudotime between a stem cell cluster and a somatic cell state.

To perform probabilistic walks between a somatic cell state and a given stem cell state, we only considered paths fully connected between the somatic cell and stem cell. Cell-cell distances were then converted to probabilities by inverting distances and normalizing the inverted score by the sum of all inverted distances for cells in the same neighborhood. Formally, cell transition probabilities were estimated using (eq. 10):

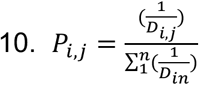

Where *P_ij_* is the transition probability between cell *i* and cell *j, D_ij_* is the Euclidean distance between cell *i* and cell *j,* and *D_in_* is distance between cell *i* and any cell sharing an edge with cell *i* (cell *n*). We then performed 1,000 probabilistic walks with a maximum step number of 1,000 between the start and end cells for each dedifferentiation trajectory conditioned by increasing pseudotime and cell state transition probabilities.

### Phytohormone dedifferentiation-bias logistic regression

To determine if phytohormone treatments were associated with increased differentiation in the various trajectories, we modeled hormone treatment status as a function of consensus pseudotime using logistic regression for each probabilistic walk (eq. 11).

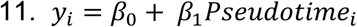

Where *y_i_* represents a numeric vector of hormone treatment status [0,1] of cell *i*, and *Pseudotime_i_* represents the consensus pseudotime value of cell *i.* Significant effects of pseudotime on hormone status was determined by empirical false discovery analysis by permuting cell pseudotime and hormone status labels, rerunning logistic regression, and collecting the null coefficient statistic from 100 permutations. Empirical *P-*values were estimated by standardizing observed statistics relative to the null distributions. Empirical *P-*values were adjusted for multiple testing across all probabilistic walks and trajectories using *p.adjust* with non-default settings (method=”fdr”). Fitted lines for each probabilistic walk were collected using the base R function, *predict*, with the model object and non-default settings (type=”response”). The number of significant walks was determined using an FDR cut-off of 0.05 and used to estimate the proportion of walks associated with increasing, decreasing, or no association with phytohormone treatments as a function of consensus pseudotime.

### Dedifferentiation velocity

The time-resolved nature of these data provided a unique opportunity to quantify dedifferentiation velocity across each dedifferentiation trajectory. To this end, we developed generalized additive mixed effect models to model dedifferentiation rates as a function of hormone treatment status and real time. Technical variation and pseudoreplication bias was controlled by including the library preparation labels as a mixed effect. Effects were estimated with REML. The formal model is presented in eq. 12.

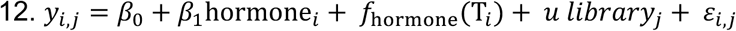

Where *y_i_*_j_ represents consensus pseudotime for cell *i* in replicate *j,* hormone*_i_* represents hormone treatment status, *f*_hormone_(T*_i_*) represents a treatment-specific smooth function over time, and *library*_j_ represents a random intercept term for library *j.* Dedifferentiation velocity was estimated from these data as the first order derivative for hormone and non-hormone treated samples, independently, using the function, *derivatives*, from the R package *gratia* (v0.11.2) with non-default settings (type=”central”).

### Reverse-pseudotime associated dynamic gene expression

To identify differentially expressed genes along each dedifferentiation trajectory, we subset the log-normalized cell x gene expression matrix to cells traversed along at least one probabilistic walk. After sorting the gene expression matrix by increasing pseudotime, we performed regression using generalized additive models for each gene using a similar strategy as for time-resolved differential expression due to hormone treatment. Specifically, dedifferentiation trajectory-associated genes were identified by comparing a full (eq. 13) and reduced (eq. 14) model lacking a b-spline smoothing term with a generalized likelihood ratio test:

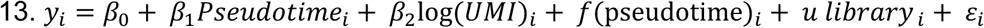

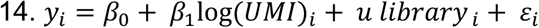

False discovery rates were estimated from the *P-*values of the generalized likelihood ratio test using the Benjamini-Hochberg method with the R function, *p.adjust* with non-default parameters (method=”fdr”). Genes with FDR < 0.01 were considered significantly associated with dynamic expression patterns along the dedifferentiation trajectory. To ensure that dedifferentiation-associated genes were free of spurious associations, we repeated the generalized likelihood ratio tests from above for each gene and trajectory after randomizing the pseudotime ordering. All genes with FDR < 0.01 in the permuted pseudotime model in the same trajectory were excluded.

### Dynamic time-warping alignment of dedifferentiation trajectories

The R package, *cellAlign* (v0.1.0), was used to identify global and single-gene expression alignments with dynamic time-warping among unrelated trajectories^75^. Unrelated trajectories were defined as pairs of trajectories with distinct somatic origins and distinct stem cell cluster fates. For each trajectory, cells traversed in at least one random walk were used to subset the log-normalized cell x gene expression matrix, followed by sorting the columns (cells) by pseudotime. Genes expressed in at least one cell were retained. Smoothed and scaled expression levels for each gene was estimated using the functions, *interWeights* and *scaleInterpolate,* from *cellAlign* with non-default parameters (trajCond=trajectory, winSz=0.1, numPts=300) and default parameters, respectively. Distance matrices were then calculated for all pairwise trajectory combinations with the function, *calcDistMat* with non-default arguments (traj.expr[[i]], traj.expr[[j]], dist.method=”Euclidean”) using the intersection of expressed genes between trajectories. Global alignments were estimated using the *globalAlign* function with non-default parameters (distMat, scores=list(query=traj[[i]], ref=traj[[j]], sigCalc=F, numPerm=100)). Single gene alignments were conducted by iterating over the shared gene list using the *globalAlign* function with non-default settings (traj.expr[[i]][x,], traj.expr[[j]][x,], scores=list(query=traj[[i]], ref=traj[[j]], sigCalc=T, numPerm=100)). An empirical FDR approach was taken to identify thresholds for considering genes as significantly aligned for each pairwise comparison. Briefly, we re-ran single gene alignments, but instead of using the canonical pseudotime estimates for each trajectory, we randomized cell-pseudotime labels. As gene expression patterns are randomized for both trajectories, the resulting distribution of normalized distances scores reflects a null distribution of alignments. Using the null distribution for each pairwise comparison, we then filtered genes from the canonical alignments with normalized distances (ranging from 0 to 0.5) less than or equal to the lesser of the bottom 1% (eFDR < 0.01) of null normalized distances or an absolute cut-off of 0.05. Residuals for aligned genes (passing eFDR thresholds) was estimated as the absolute difference in the global alignment position (trajectory 1 – trajectory 2) divided by the square root of 2. The average alignment residual in the early, middle, and late stages of differentiation were estimated by summing residuals over the first third, middle third, and last third of the aligned trajectories. K-means clustering was conducted by first averaging normalized gene expression patterns (ranging from 0-1) across all trajectories and standardizing averaged expression patterns for each gene. The optimal *k-*means clusters was determined by a combination of the elbow, silhouette, and gap statistic methods. Shannon entropy values for each gene and reverse pseudotime point were determined by converting expression values across trajectories into a probability distribution and taking the negative sum of the expression probabilities multiplied by the log_2_ of the probability distribution. Gene set enrichment analysis was conducted by estimating the distance of each aligned gene to all k-means centroids, converting distance estimates to negative scores by multiplying by negative one, and standardizing adjusted distance metrics across k-means clusters, which can be interpreted as gene enrichment within a specific cluster. Gene set enrichment analysis was then applied using the enrichment scores across all aligned genes to each k-means cluster using *fgsea* with non-default parameters (minSize = 5, maxSize = 400, nPermSimple = 10000).

### Identification of early dedifferentiation regulators

To identify putative regulators of early dedifferentiation onset, we generated dGRNs for each dedifferentiation trajectory. Specifically, cells from all probabilistic walks per trajectory were collapsed, taking the union of cell identifiers. The merged set of cell IDs were ordered by reverse pseudotime consensus values and then used to subset and sort the cell x log-normalized expression matrix. Activation timing, *A*, for gene *i* was then estimated using the center of mass (eq. 15):

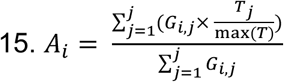

Where *G_ij_* is the log-normalized expression level of gene *i* in cell *j* and *T_j_* is the pseudotime ranking of cell *j.* Genes with putative regulator function were identified on the basis of GO terms for transcription factors (GO:0003700, GO:0140110, GO:0001228, GO:0001227, GO:0000976, GO:0000981), chromatin remodelers (GO:0006338, GO:0140657, GO:0016570, GO:0016573, GO:0016571, GO:0006306, GO:0003682), phosphoregulators (GO:0016301, GO:0004672, GO:0004674, GO:0004713, GO:0016791, GO:0004721), and signal transduction (GO:0007165, GO:0004871, GO:0005102). Regulators in the top 20% of activation timing expressed in at least two independent trajectories were retained. The filtered set of regulators were used to construct dynamic GRNs for each dedifferentiation trajectory with a multi-lagged approach. Briefly, the cell x log-normalized expression matrix was subset and ordered by reverse pseudotime consensus values, filtered regulators, and genes with non-zero expression in at least 5 cells. Expression levels for each gene were averaged within 100 bins based on cell reverse pseudotime consensus and then standardized across bins (mean subtracted and divided by the standard deviation). Regulator-target correlation matrices were then calculated for a range of pseudotime bin delays (1, 2, 3, 5, 10, and 20 bins), enabling estimates of time-delayed effects of regulator expression across progressively distant pseudotime bins. Regulator-target correlations were weighted by the normalized exponential decay across all pseudotime lags, resulting in a weighted average regulator-target correlation across multiple lag estimates, using eq 16.

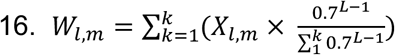

Where *X_lm_* is the Pearson’s correlation coefficient between regulator gene *l* at pseudotime bin *u* and target gene *m* expression levels at pseudotime *u* + lag and *L* is the vector of bin lagged values (1, 2, 3, 5, 10, and 20). Weighted Pearson’s correlations above 0.8 and below -0.8 were retained, all other regulator-target edges were dropped.

The filtered set of regulator-target pairs were then compared across all dedifferentiation trajectories. First, only regulator-target pairs with an absolute weighted correlation average greater than 0.8, supported by at least three trajectories were retained, and at least two trajectories initiated in distinct somatic cell type origins were retained. The number of concordant regulator target genes was estimated by intersecting regulator-target pairs across all trajectories and counting the number of target genes per regulator consistently observed across multiple trajectories (i.e. conserved regulator-target pairs across distinct trajectories). The number of conserved target genes, average activation timing, and fraction of trajectories in which a regulator is expressed were used to derive a dedifferentiation initiation score for each regulator (eq. 17):

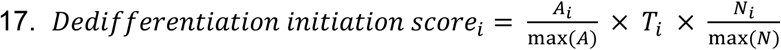

Where *A_i_* is the average activation timing of gene *i, T_i_* is the proportion of trajectories where gene *i* is expressed, and *N_i_* is the number of conserved target genes for regulator gene *i*.

### Transcriptional divergence from somatic origin cell types

Generalized additive models were used to identify the timing and kinetic of dedifferentiation and stem cell heterogeneity. Briefly, the union of cell identifiers from each dedifferentiation trajectory were collected from 1,000 probabilistic walks, sorted by reverse pseudotime consensus, and used to subset the cell x log-normalized gene expression matrix. To quantify the number of differentially expressed genes at discrete reverse pseudotime points with respect to the initial somatic state, we modeled gene expression as a smooth function using generalized additive models. For each gene *g,* expression values across cells were fit independently using eq. 18:

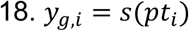

Where *y_g,I_* denotes the log-normalized expression levels of gene *g* in cell *i,* and *s()* is a penalized regression spline with a fixed number (*k*=6) of basis functions. The models were fit with a Gaussian error distribution. To estimate the number of gene expression changes at any given timepoint, the reference timepoint *T_0_* was defined as the minimum reverse pseudotime consensus value after rescaling each trajectory from 0 to 1, where 0 represents somatic cell states and 1 represents stem-cell like identities. We then evaluated gene expression changes on a dense grid (100 evenly spaced points) of rescaled pseudotime between 0 and 1. The difference between predicted expression levels between each grid point *T_i_* and the reference time point *T_0_* was estimated across all genes using the linear predictor matrix of the fitted generalized additive model. To estimate statistical significance of temporal deltas, we estimated uncertainty using the covariance matrix of the model coefficients. Specifically, the variance of each timing contrast was computed using eq. 19:

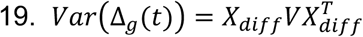

Where *X_diff_* represents the difference between the design matrices at *T_i_* and *T_0_*, and *V* is the coefficient covariance matrix. Wald statistics were then estimated using the standard errors from the delta variance via eq. 20:

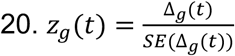

Two-sided *P-*values were estimated assuming asymptotic normality. FDR scores were estimated from the raw *P-*values using the R function, *p.adjust* with non-default parameters (method=”BH”) for each temporal pairwise comparison. Genes were defined as differentially expressed if FDR < 0.05 and the absolute effect size (Δ_g_(*t*)) was greater than 0.25. Differential expression velocity within each trajectory was estimated using the first derivative of a generalized additive model fit by the number of differentially expressed genes at each timepoint using the function, *derivatives* from the R package, *gratia* with non-default parameters (fit, select = “s(t)”, n=100).

### Use of generative AI

Claude (Opus 4.3) was used to optimize and annotate code related to TF and nucleosome footprinting and was extensively evaluated with human expert oversight to ensure reproducibility and biological relevance. Prompts and Claude summaries are directory available in the GitHub repository (Marand-Lab/Arabidopsis_Dedifferentiation_scRNAseq_Atlas) in the subdirectory named /08_protoplast_ATAC_analysis.

## DATA AVAILABILITY

Raw and processed data associated with this study, including a fully specified Seurat object complete with cell annotations, dedifferentiation scores, reverse pseudotime, and various embeddings, can be found freely and publicly available at NCBI GEO following peer review.

## CODE AVAILABILITY

Custom and publicly available code used in this study has been archived through Zenodo^76^ and can be found freely available on GitHub at the following repository URL: https://github.com/Marand-Lab/Arabidopsis_Dedifferentiation_scRNAseq_Atlas.

## ACKNOWLEDGEMENTS

We wish to thank Dr. Libo Shan, Dr. Ping He, Dr. Robert J. Schmitz, and Dr. Tyler Huycke for providing valuable feedback on the manuscript. We are also grateful for the Advanced Research Computing at the University of Michigan, Ann Arbor for providing computational resources and technical support. This work was supported by the United States National Science Foundation (IOS 2314549, IOS 2424273) and USDA National Institute of Food and Agriculture, HATCH project VA-160262, and Multistate S-009 project VA-136423 to B.O.R.B, the Momental Foundation and Michigan Pioneer Fellows Program to F.G.C., and the National Institutes of Health (R00GM144742) and start-up funds from the University of Michigan to A.P.M.

## CONTRIBUTIONS

A.P.M. conceived and designed the experiments. A.P.M. and B.O.R.B. supervised the study. Y.C., M.C.G., K.H.R., L.V., and K.M.R. performed experiments. F.G.C., L.J., M.C.G., K.H.R., R.R., L.V., B.O.R.B., and A.P.M. analyzed the data. A.P.M. wrote the manuscript with input from all authors. All authors approved the final manuscript.

## Supplemental Figures

**Figure S1:**
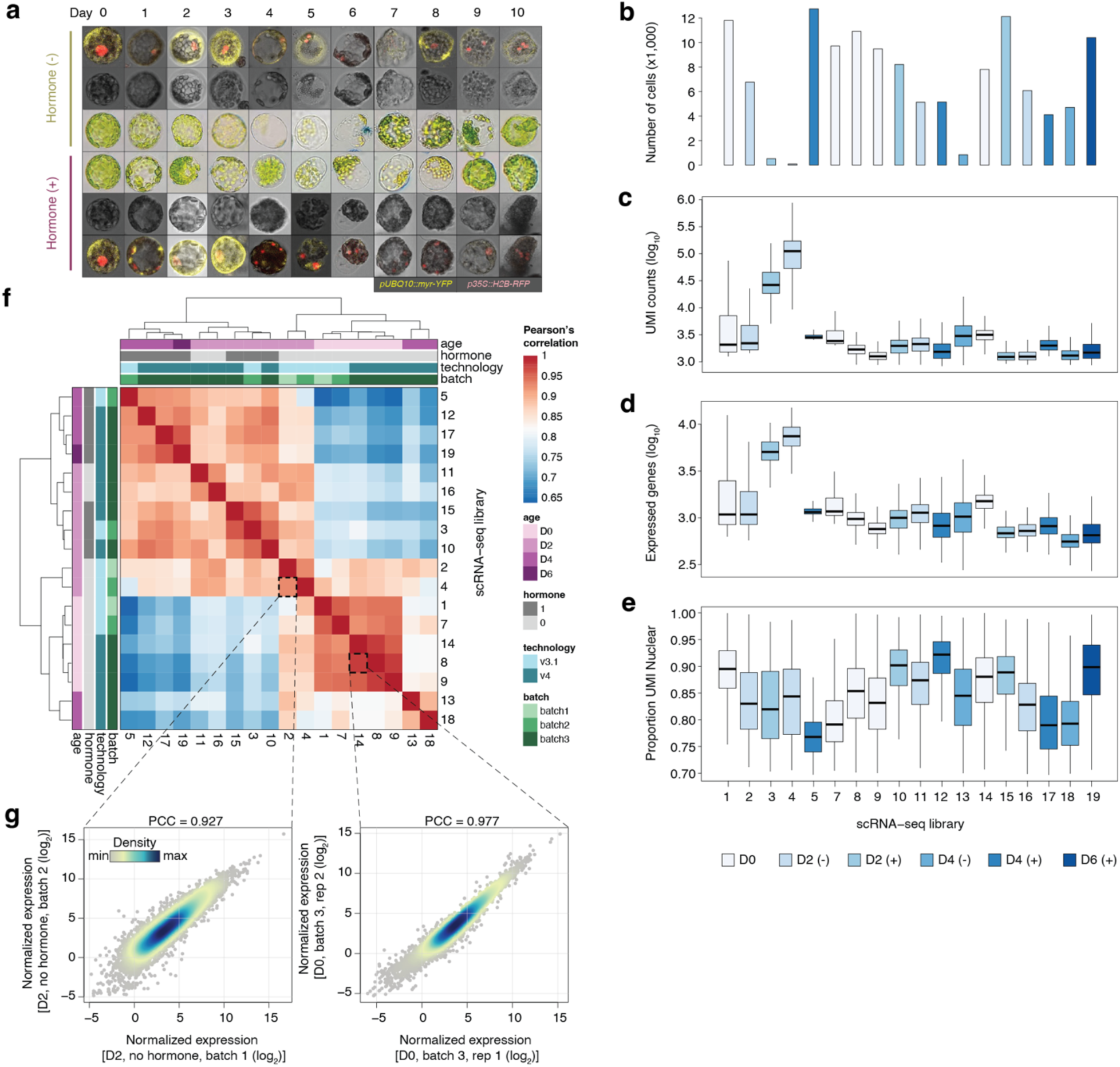
Single-cell RNA-seq quality control. (**a**) Representative brightfield, light (middle), and merged confocal micrographs (top and bottom) of a Day 0 nascent protoplast (the same protoplast is shown for both hormone and non-hormone treated for consistency) and Days 1-10 protoplast samples treated with and without 2,4-D and 6-BA. Red fluorescence indicates *p35S::H2B-RFP* expression and yellow indicates *pUBQ10::myr-YFP* expression. (**b**) Bar plot illustrating the number of QC filtered cells in each library. (**c**) Boxplots of the log_10_ transformed UMI counts across cells in each library. (**d**) Boxplots of the log_10_ transformed number of expressed genes across cells in each library. (**e**) Distributions of the proportion of nuclear-derived UMIs across cells in each library. (**f**) Pearson’s correlation coefficient matrix of all scRNA-seq libraries treated as bulk samples. Columns and rows are annotated with protoplast duration age, hormone treatment status, 10X Genomics scRNA-seq technology version, and batch. (**g**) Examples of genome-wide log_2_ normalized expression values between replicates from different batches (left) and the same batch (right).

**Figure S2:**
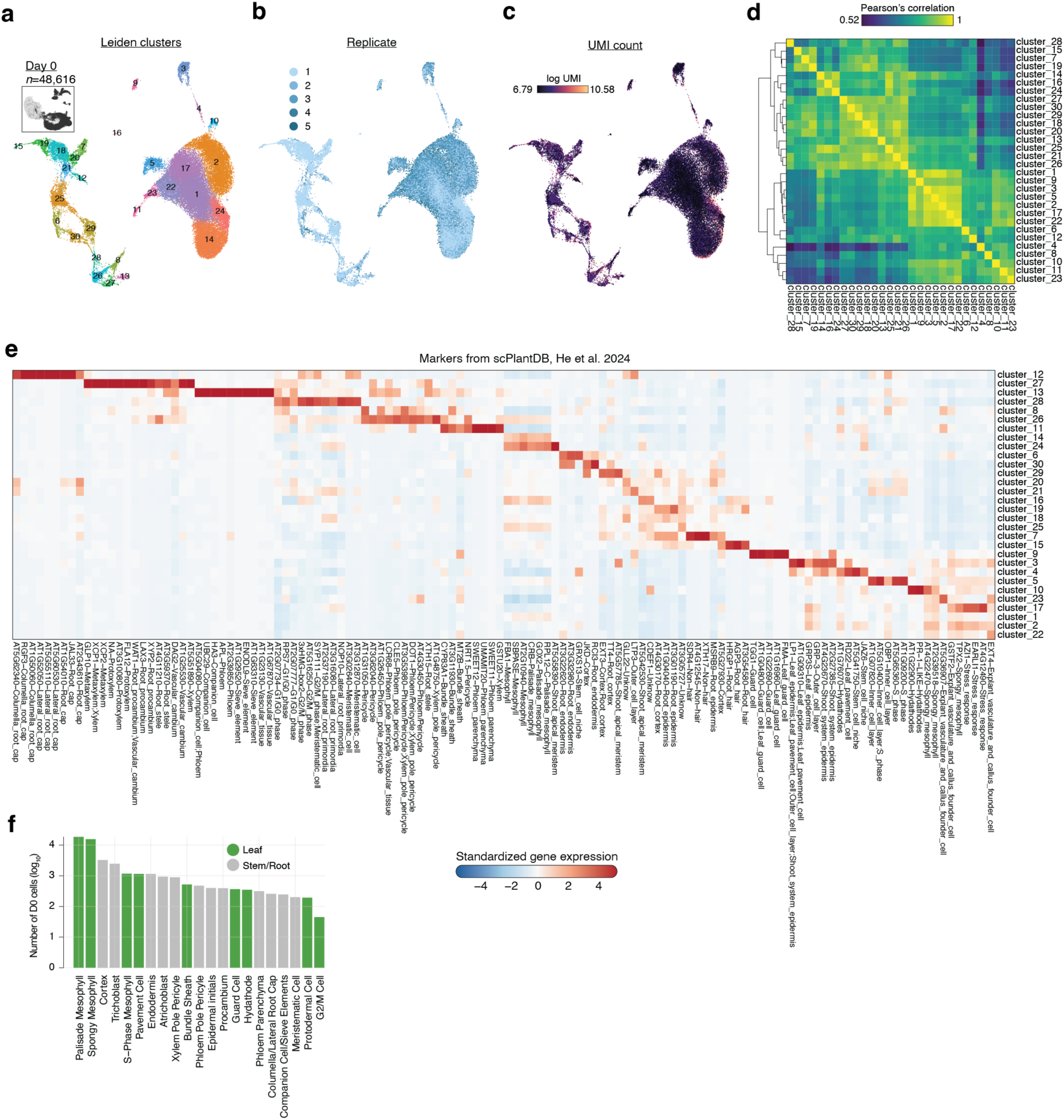
Day 0 protoplast clustering and cell type annotation. (**a**) Leiden cluster assignments from the shared nearest neighbor graph visualized on the UMAP embedding. (**b**) UMAP embedding of cells colored by replicate. (**c**) UMAP embedding of cells colored by log transformed UMI counts. (**d**) Pearson’s correlation coefficient heatmap of cluster-averaged normalized gene expression values. (**e**) Heatmap of standardized gene expression values (Z-scores) across clusters for the curated set of cell type marker genes. (**f**) Barplot illustrating Day 0 cell counts for each annotated cell type, colored by the tissue where the cell type occurs most frequently.

**Figure S3:**
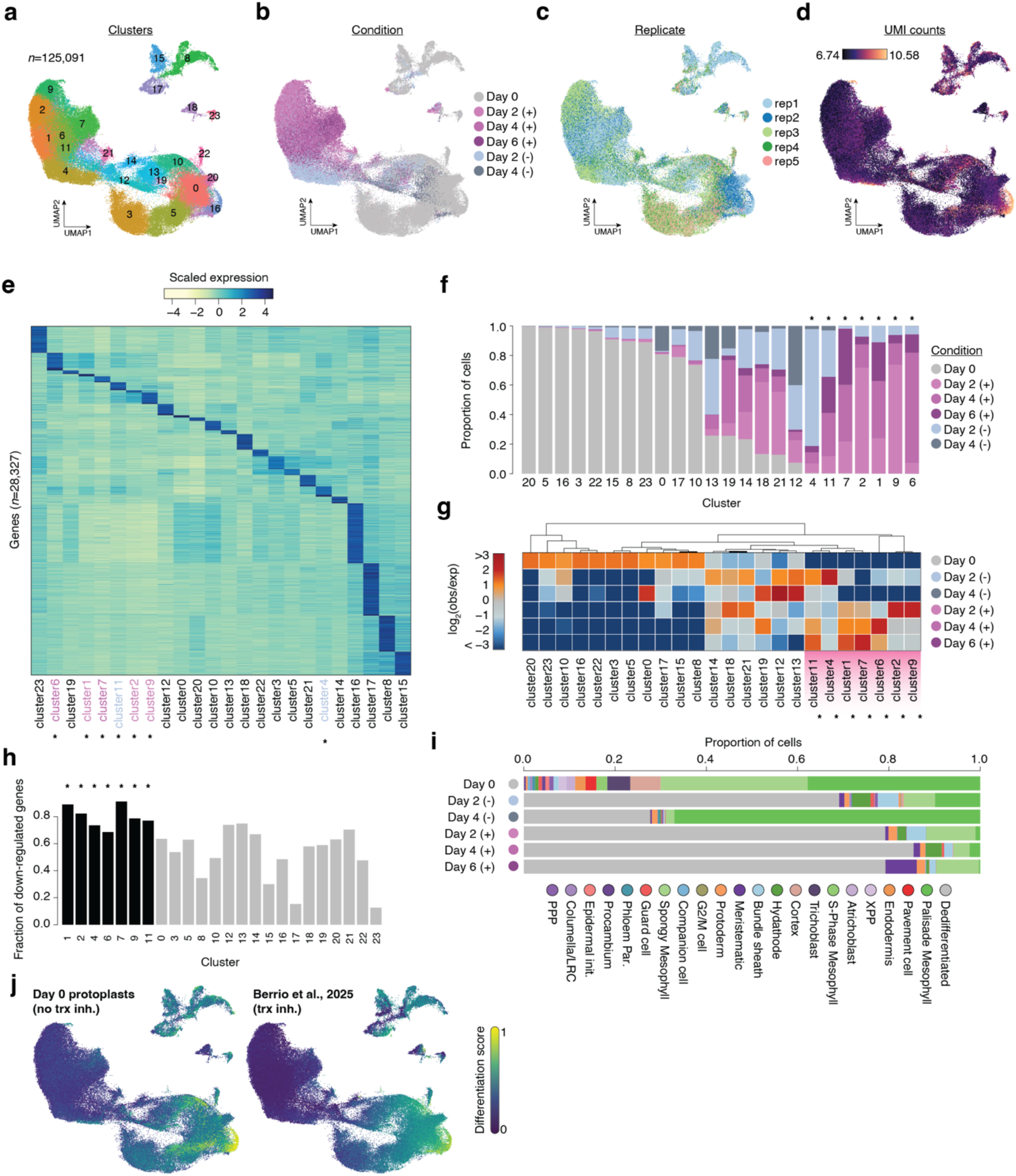
Quality control for the integrated scRNA-seq protoplast experiment. (**a**) UMAP embedding of all scRNA-seq protoplasts colored by cluster assignment. (**b**) UMAP embedding of all scRNA-seq protoplasts colored by experimental condition. (**c**) UMAP embedding of all scRNA-seq protoplasts colored by replicate. (**d**) UMAP embedding of log transformed UMI counts for all scRNA-seq protoplasts. (**e**) Heatmap of standardized gene expression values (Z-scores) across clusters from the fully integrated data set. Clusters with significant (FDR < 0.05 & log_2_ fold-change < -3) depletion of Day 0 are indicated by asterisks. (**f**) Bar plot of the proportion of cells derived from different experimental conditions within each cluster. Clusters with significant (FDR < 0.05 & log_2_ fold-change < -3) depletion of Day 0 are indicated by asterisks. (**g**) Heatmap of the log_2_ observed over expected cell frequencies partitioned by cluster. Clusters with significant (FDR < 0.05 & log_2_ fold-change < -3) depletion of Day 0 are indicated by asterisks. (**h**) Bar plot of the fraction of differentially expressed genes that are downregulated in each cluster compared to all cells. Black bars indicate clusters that are statistically depleted of Day 0 cells. (**i**) Cell type proportions partitioned by experimental condition. (**j**) Comparison of differentiation scores estimated using Day 0 protoplasts (left; this study) and a published scRNA-seq atlas (right) where new transcription was inhibited with actinomycin D during cell wall digestion.

**Figure S4:**
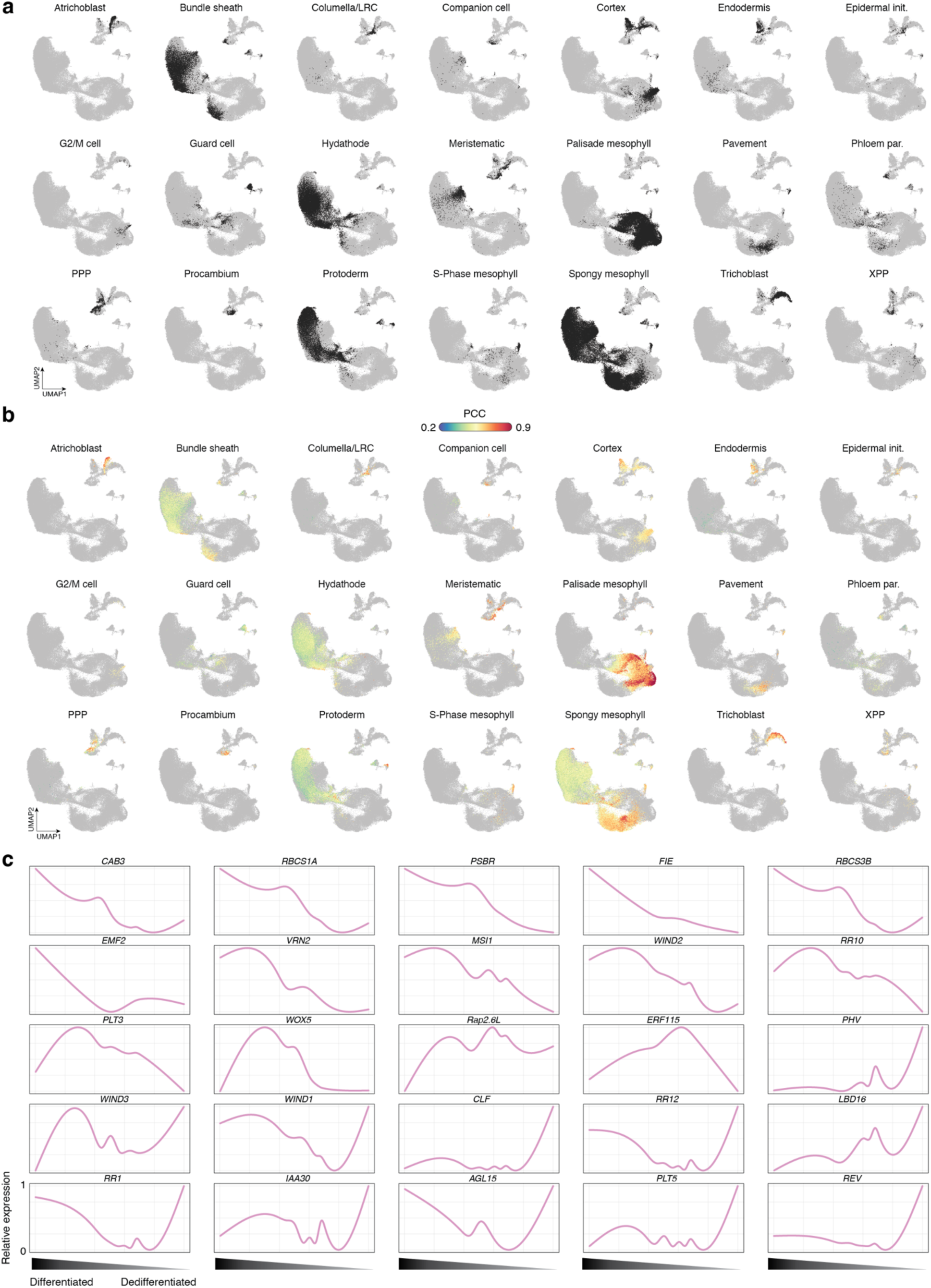
Somatic similarity and gene expression dynamics associated with dedifferentiating cells. (**a**) UMAP embedding of the best predicted cell type label from genome-wide correlations with Day 0 aggregate cell types. (**b**) UMAP embedding of the maximum Pearson’s correlation coefficient for each cell with Day 0 aggregate cell types. (**c**) Expression levels ordered by differentiation status are shown for representative marker genes.

**Figure S5:**
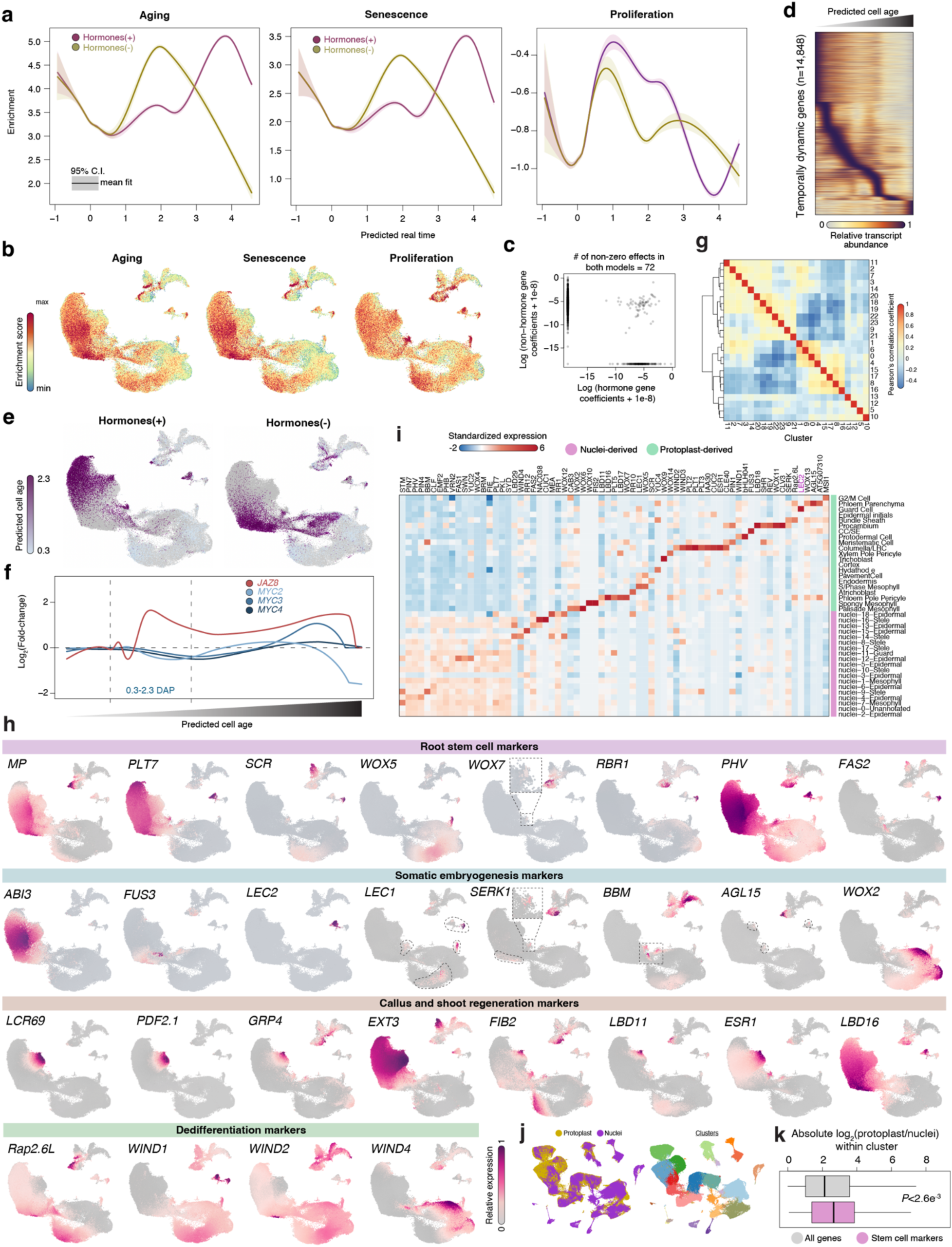
Phytohormone effects and marker gene expression diversity in dedifferentiating protoplasts. (**a**) Gene set enrichment profiles across time partitioned by hormone and non-hormone treated samples. (**b**) Per cell gene set enrichment scores illustrated on the integrated UMAP embedding. **(c)** Comparison of LASSO regularized regression coefficients (gene effects) from the cell age models using the transcriptomes of non-hormone (y-axis) and hormone-treated cells (x-axis). Genes with non-zero effects in at least one of two models are shown. (**d**) Heatmap of temporally dynamic genes expression (FDR < 0.05). (**e**) LASSO regularized regression predicted cell age based on hormone (left) and non-hormone-treated (right) transcriptomes. (**f**) Log_2_ fold-change in transcript abundance between hormone-treated and non-hormone-treated samples across time for *JAZ8* and JAZ8 target genes, *MYC2/3/4*. (**g**) Heatmap visualizing pair-wise Spearman correlation coefficients of transcriptomic profiles among all clusters from the integrated data set. (**h**) UMAP embedding illustrating relative gene expression levels on the integrated data set grouped by root stem cell (purple), somatic embryogenesis (blue), callus and shoot regeneration (light brown), and dedifferentiation (green) markers. (**i**) Expression patterns of stem cell marker genes across 11-day old seedling protoplast cells (light green clusters) and 12-day old seedling nuclei (purple clusters). (**j**) Left, UMAP embedding indicating integration of protoplast (gold points) and nuclei (purple points) derived from Arabidopsis seedlings. Right, UMAP embedding indicating Leiden cluster assignments. (**k**) Distribution of absolute log2 fold-changes between protoplast and nuclei observations across integrated clusters for stem cell markers (pink) and all genes (grey).

**Figure S6:**
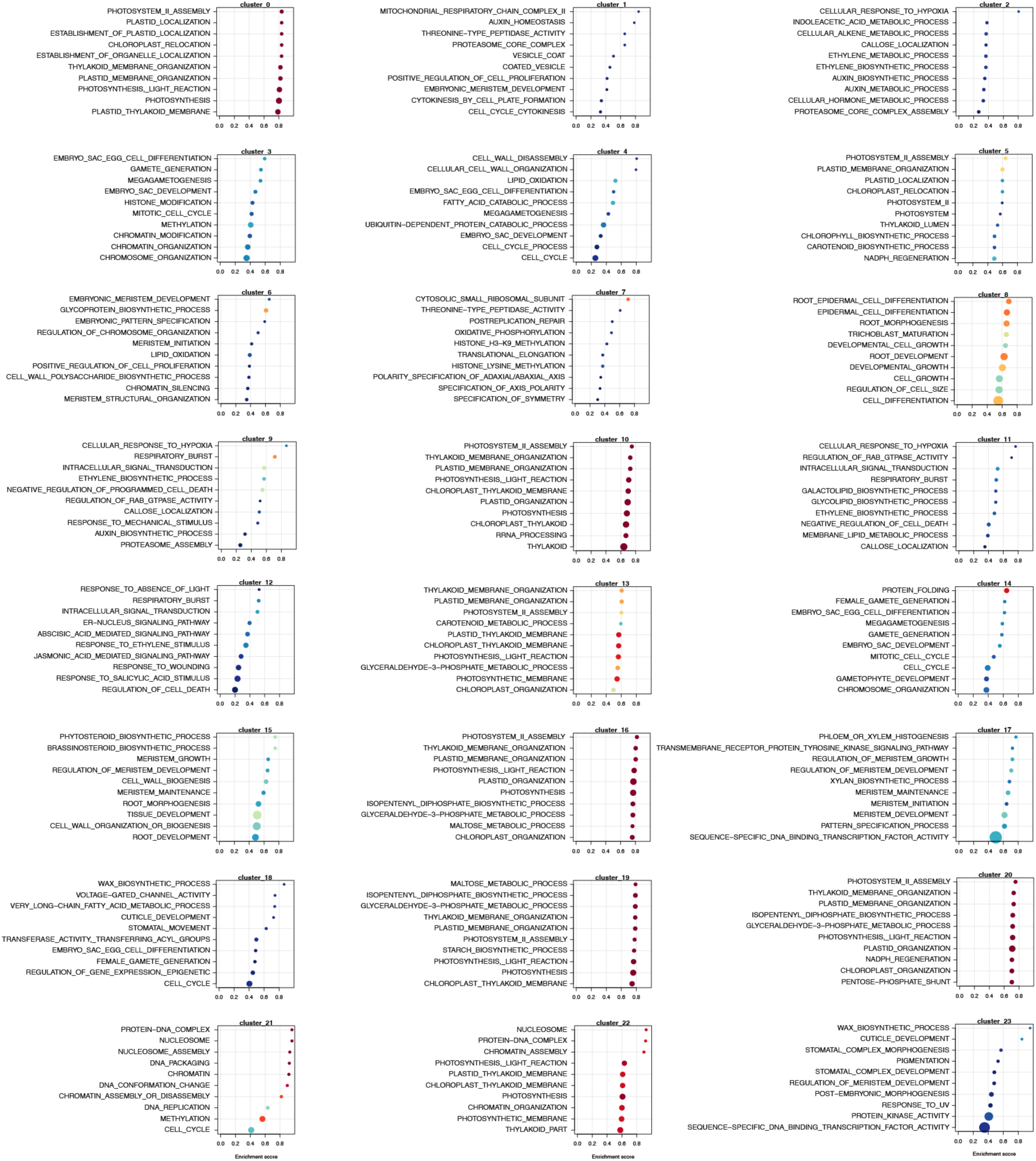
Gene set enrichment analysis. Ten representative gene annotation enrichment metrics for each cluster from the integrated data set.

**Figure S7:**
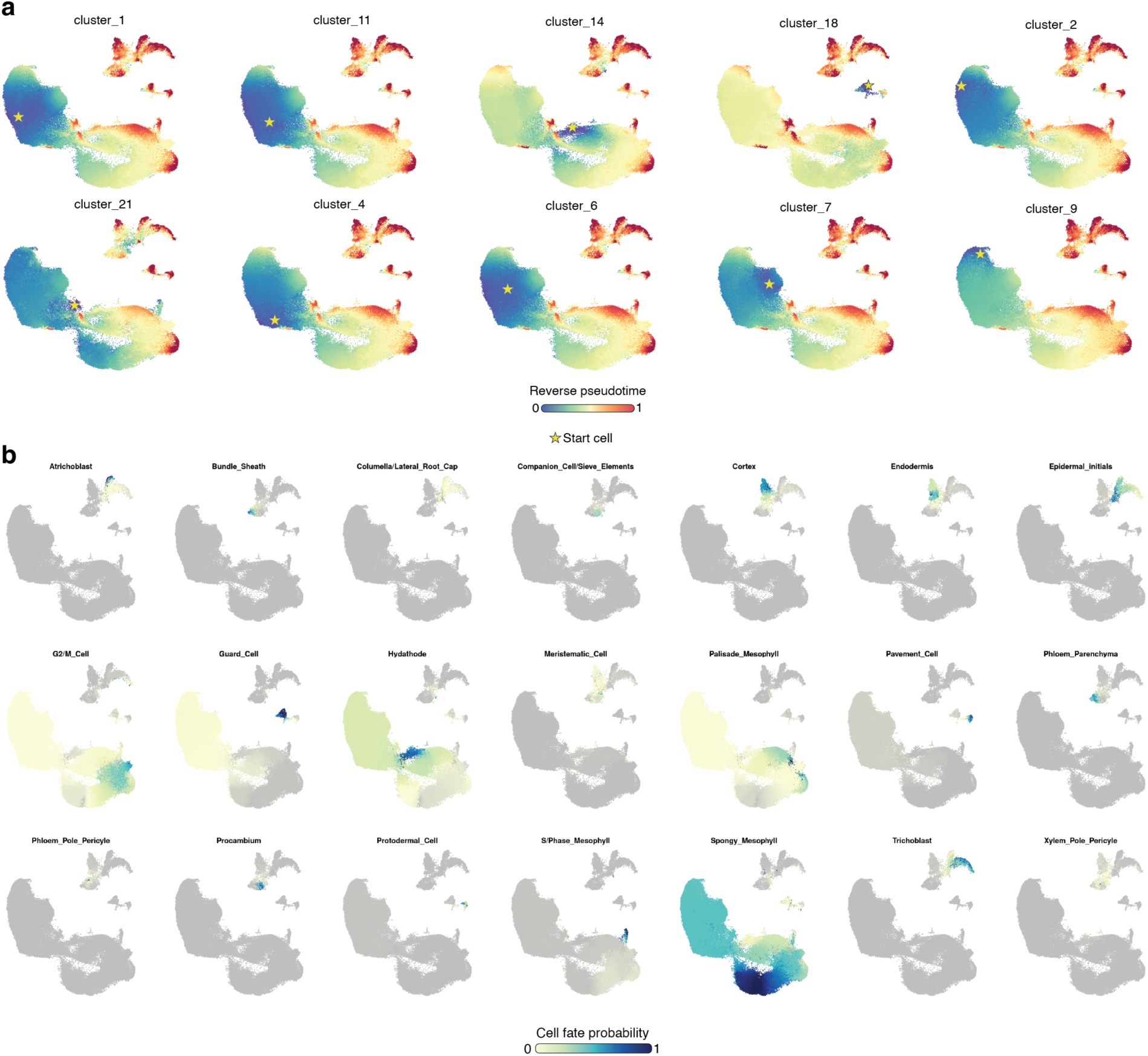
Evaluation of reverse pseudotime and cell fate probabilities across diverse initial somatic cells and stem cell-like fates. (**a**) UMAP visualization of reverse pseudotime scores from modeling each candidate cluster as a stem cell initial. Gold start indicates trajectory start cell. (**b**) Cell fate probabilities for each somatic cell type.

**Figure S8:**
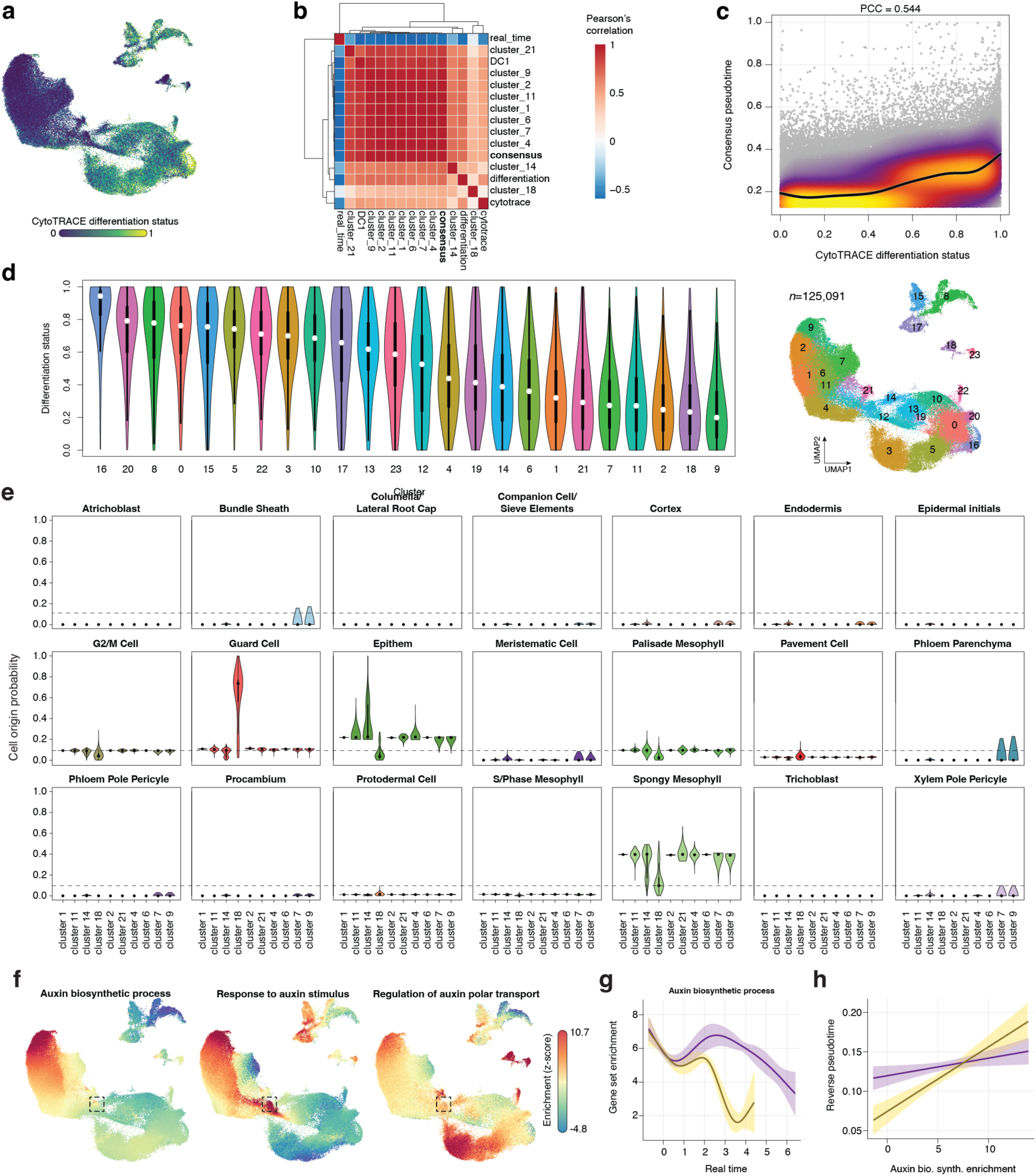
Orthogonal metrics support diffusion-based pseudotime. (**a**) UMAP embedding visualization of CytoTRACE-based differentiation status scores. (**b**) Heatmap indicating the pairwise correlation among reverse pseudotime estimates from the 10 trajectories, cytotrace differentiation scores, diffusion component 1, and real time. (**c**) Density scatter plot comparing differentiation status from CytoTRACE (x-axis) with the consensus (minimum) reverse pseudotime estimates from Palantir (y-axis). **(d)** Violin plots summarizing differentiation status distribution within each cluster from the integrated embedding (left). The UMAP embedding with cluster assignment is visualized for convenience (right). (**e**) Distribution of cell fate probabilities for the 10 stem cell clusters across all observed terminally differentiated cell types. (**f**) UMAP embedding of standardized (z-score) gene set enrichment scores per cell for auxin biosynthetic process (left), response to auxin stimulus (middle), and regulation of auxin polar transport (right). (**g**) Gene set enrichment dynamics across real time for hormone (purple) and non-hormone (yellow) epithem and epithem-derived protoplasts. Regression fit and 95% confidence interval is indicated by the solid line and shaded polygon, respectively. (**h**) Effect of auxin biosynthesis gene expression on reverse pseudotime estimates partitioned by hormone (purple) and non-hormone (yellow) treated protoplasts. Regression fit and 95% confidence interval is indicated by the solid line and shaded polygon, respectively.

**Figure S9:**
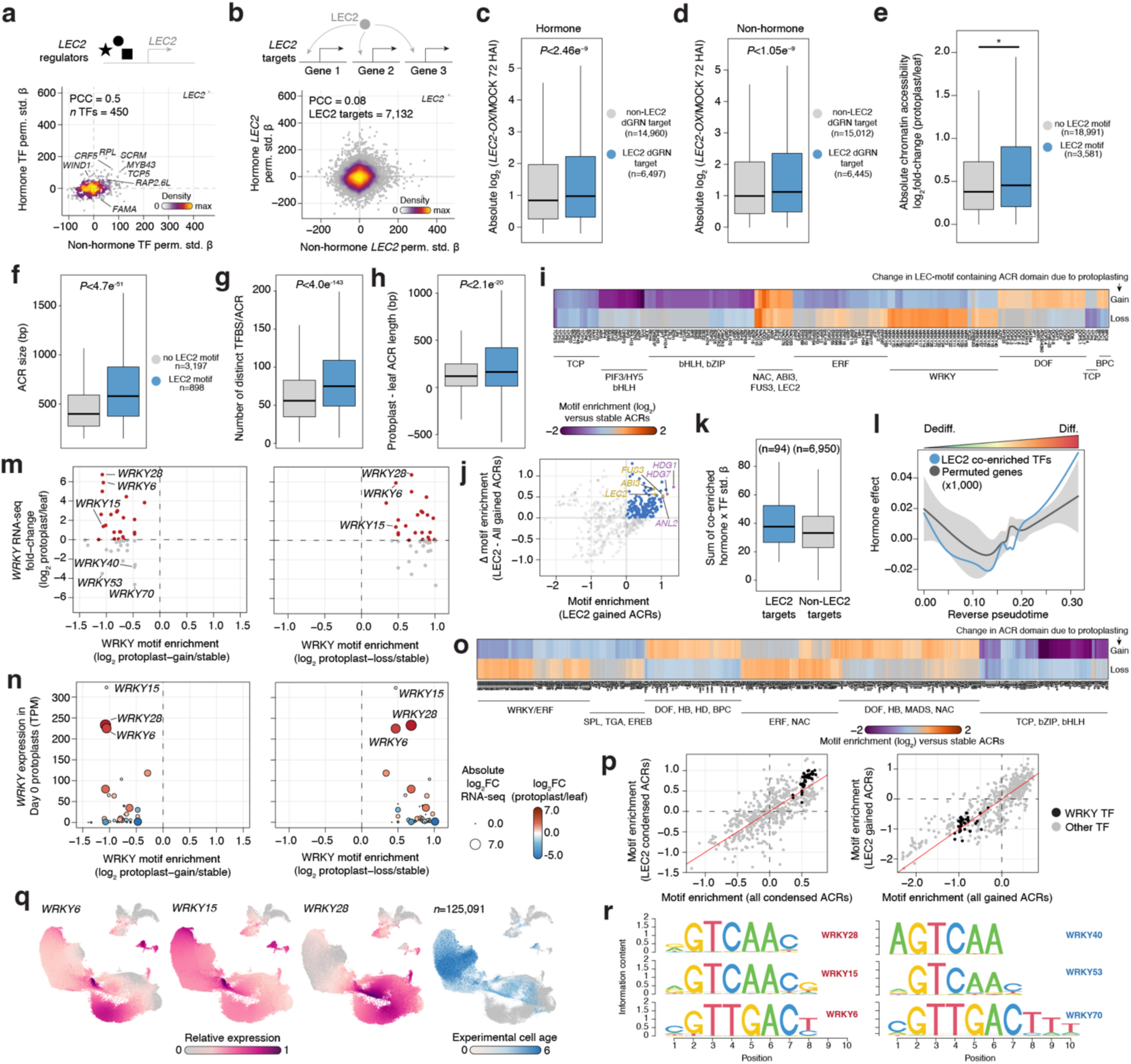
Dynamic gene regulatory networks underlying dedifferentiation. (**a**) Density scatterplot of permutation standardized TF effects on *LEC2* expression from dynamic gene regulatory networks constructed from hormone (y-axis) and non-hormone (x-axis) treated protoplasts. (**b**) Density scatterplot of permutation standardized *LEC2* effects on *LEC2* target gene expression from dynamic gene regulatory networks constructed from hormone (y-axis) and non-hormone (x-axis) treated protoplasts. (**c**) Distribution of absolute log_2_ fold-change (*LEC2* over-expression versus mock) after 72h of induction for LEC2 and non-LEC2 targets in hormone-treated protoplasts. (**d**) Distribution of absolute log_2_ fold-change (*LEC2* over-expression versus mock) after 72h of induction for LEC2 and non-LEC2 targets in non-hormone-treated protoplasts. (**e**) Distribution of absolute chromatin accessibility log2 fold-change between protoplasts and intact leaves partitioned by ACRs with (blue) and without (grey) LEC2 DNA motifs. (**f**) Distribution of ACR size (bp) between ACRs with (blue) and without (grey) LEC2 motifs. (**g**) Distribution of distinct TF motif counts per ACR for ACRs with (blue) and without (grey) LEC2 motifs. (**h**) Distribution of ACR size change (protoplast ACR length – intact leaf ACR length) upon protoplasting for ACRs with (blue) and without (grey) LEC2 motifs. (**i**) Heatmap of TF motif enrichment (log_2_ fold-change) in LEC2 motif-containing ACRs unique to protoplasts (gain) or intact leaf (loss) relative to stable (accessible in both protoplasts and intact leaves) ACRs. (**j**) Scatter plot of LEC2 co-motif enrichment in protoplast-unique ACRs (LEC2 gained ACRs) relative to the difference in motif enrichment between LEC2-containing ACRs and all protoplast-specific ACRs (agnostic to LEC2 motifs). Blue dots indicate LEC2 co-enriched TF motifs. (**k**) Distribution of summed hormone x TF standardized effects of LEC2 co-enriched TFs at LEC2 targets (blue) and non-LEC2 target genes (grey). (**l**) Reverse pseudotime resolved hormone effect (hormone – non-hormone gene expression) for LEC2 co-enriched TFs (blue) and 1,000 permutations of the same number of random expressed genes. The light grey polygon represents the 99% quantile of permuted values. Dark grey indicates the mean hormone effect of permuted genes. (**m**) Scatter plot of *WRKY* log_2_ fold-change in expression between intact leaves and protoplasts (y-axis) and the relative change in cognate WRKY motif enrichment for protoplast-gained ACRs (left) and ACRs lost in protoplasts (right). (**n**) Scatter plot of *WRKY* expression levels (y-axis) relative to WRKY motif enrichment in protoplast-gained ACRs (left) and ACRs absent in protoplasts (right). The size of each point represents the absolute log_2_ fold-change in expression between protoplast and intact leaves. The color of each point indicates the log_2_ fold-change direction (blue, downregulated; red, upregulated) between protoplasts and intact leaves. (**o**) Heatmap of TF motif enrichment for all ACRs unique to protoplasts (gain) or intact leaves (loss) relative to stable (accessible in both protoplasts and intact leaves) ACRs. (**p**) Scatter plot comparison of motif enrichment (log_2_ relative to stable ACRs) for ACRs containing LEC2 motifs (y-axis) and all ACRs agnostic to LEC2 motifs (x-axis) for protoplast-specific ACRs (right) and ACRs absent in protoplasts (left). WRKY TFs are colored in black. All other TFs are colored grey. (**q**) Relative transcript abundance of *WRKY6/16/28* across all cells in the integrated atlas and visualized on the UMAP embedding. Protoplast experimental age is illustrated with the blue color gradient. (**r**) TF motif sequences corresponding to the top upregulated (left) and downregulated *WRKY* TFs following protoplast induction.

**Figure S10:**
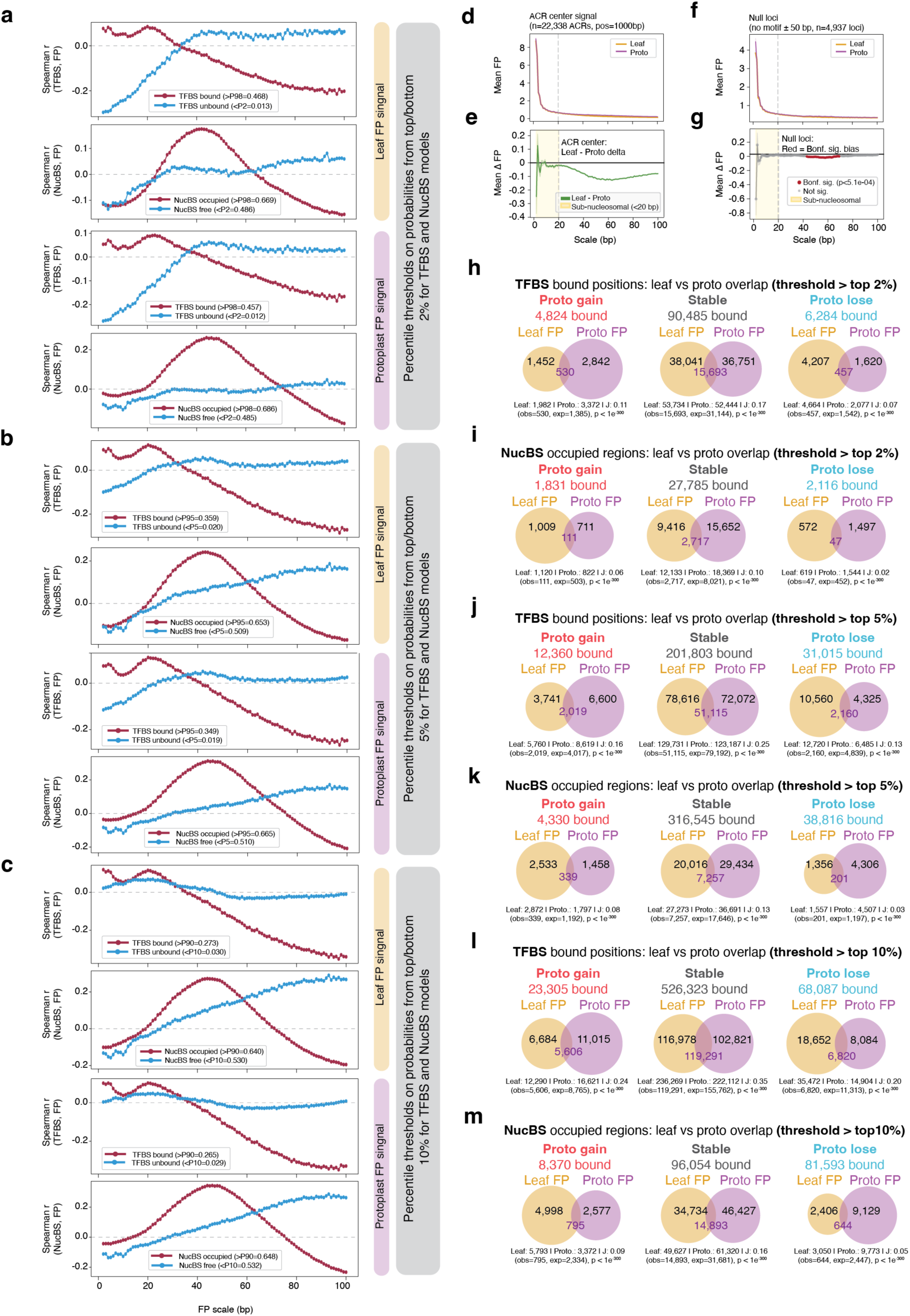
Multiscale protein footprinting in protoplast and intact leaf chromatin accessibility data. **(a**-**c)** Spearman correlations between TF/nucleosome occupancy probabilities and footprinting signal across various footprinting resolutions for intact leaf (top, top-middle) and protoplast (bottom-middle, bottom) ATAC-seq data sets using the top (red lines) and bottom (blue lines) quantiles at (**a**) 2%, (**b**) 5%, and (**c**) 10% of TF- and nucleosome-scale footprints. (**d**) Mean footprint signal at ACR centers across footprint scales for intact leaf and protoplast ATAC-seq data. (**e**) Average footprinting delta (intact leaf – protoplast) across all ACRs for various footprinting resolutions. (**f**) Average footprint score for null (no known TF motif within 50 bp) loci within ACRs for intact leaf and protoplast ATAC-seq data for various footprinting resolution. (**g**) Average footprinting delta (intact leaf – protoplast) across all ACR null positions (no known TF motif within 50bp) for various footprinting resolution. (**h**-**m**) Overlap between intact leaf and protoplast TF footprints partitioned by ACRs classified as increased chromatin accessibility in protoplasts (FDR < 0.05; left), no change in chromatin accessibility (FDR > 0.05; middle), and decreased chromatin accessibility in protoplasts (FDR < 0.05; right) for occupancy thresholds reflecting the top (**h**) 2%, (**j**) 5%, or (**l**) 10% of TF-occupied footprints. Overlap between intact leaf and protoplast nucleosome-scale footprints partitioned by ACRs classified as increased chromatin accessibility in protoplasts (FDR < 0.05; left), no change in chromatin accessibility (FDR > 0.05; middle), and decreased chromatin accessibility in protoplasts (FDR < 0.05; right) for occupancy thresholds reflecting the top (**i**) 2%, (**k**) 5%, or (**m**) 10% of nucleosome-scale footprints.

**Figure S11:**
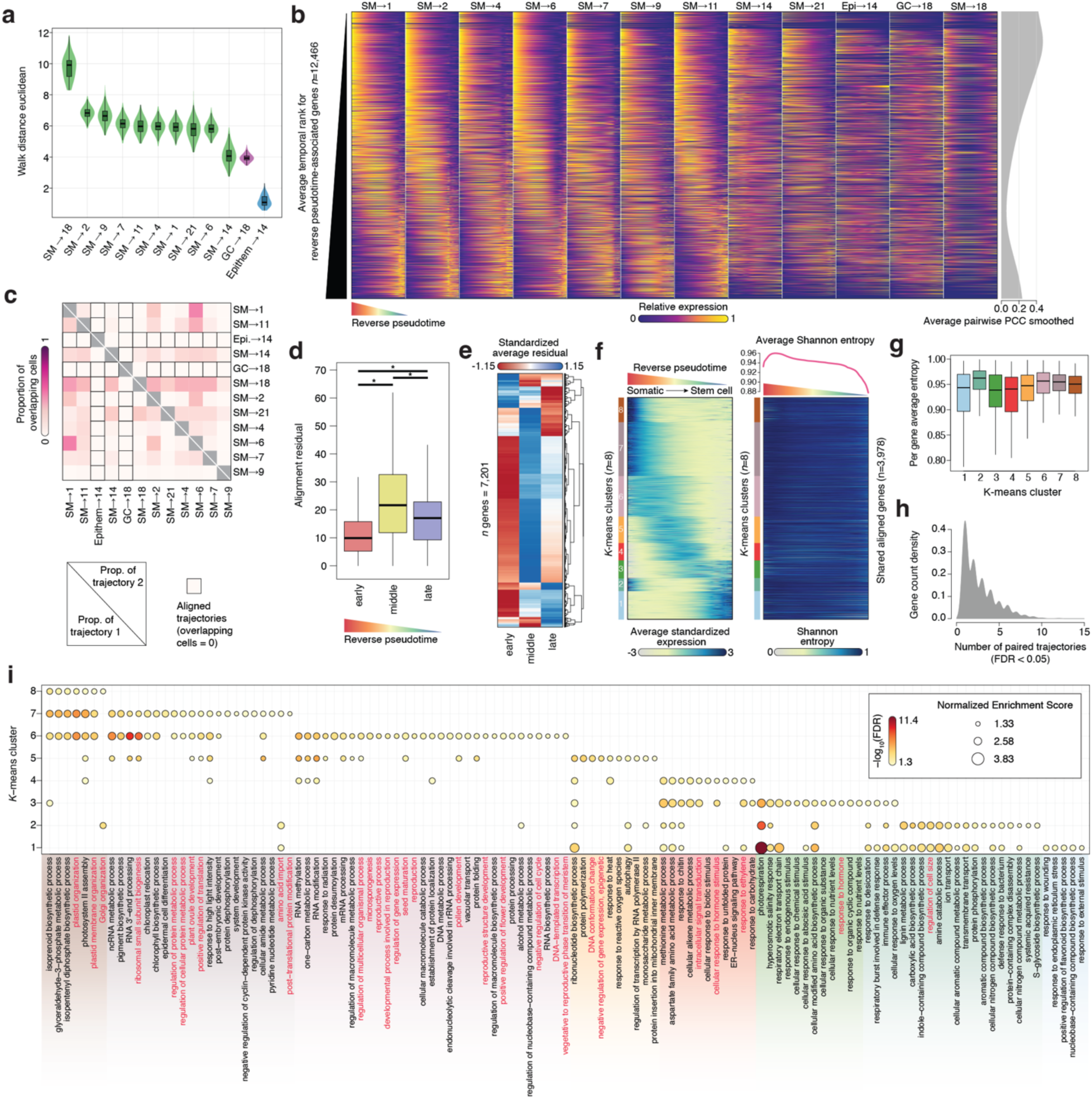
Shared and unique dedifferentiation associated gene expression patterns across distinct somatic and stem cell fate trajectories. **(a)** Distribution of probabilistic walk distances (1,000 probabilistic walks per trajectory) for each trajectory. **(b)** Heatmap illustrating dynamic gene expression patterns across the 12 dedifferentiation trajectories. Rows (genes) are ordered by their average rank across trajectories. Right, average smoothed (smoothing spline) pairwise Pearson’s correlation coefficient across genes. **(c)** Fraction of overlapping cells between trajectories using trajectory 1 (lower diagonal) and trajectory 2 (upper diagonal) as the reference set. Matrix elements with black boxes indicate unrelated alignments. **(d)** Distribution of alignment residuals across all pairwise trajectory alignments partitioned by early, middle, and late stages of dedifferentiation. **(e)** Heatmap of the standardized average residuals across trajectories for significant (eFDR) gene alignments between at least two independent trajectories. (**f**) Left; average standardized expression dynamics across the 12 trajectories. Right; Shannon entropy values throughout reverse pseudotime across the 12 trajectories. Average Shannon entropy across genes for each timepoint is illustrated via a line plot above the heatmap. (**g**) Distribution of the number of times genes were significantly aligned (eFDR < 0.05) between trajectories. (**h**) Distribution of the number of significant (FDR < 0.05) trajectory pairs in which a gene participates. (**i**) Bubble plot indicating gene set enrichment analysis results from each *k-*means cluster. Bubbles are colored by the -log_10_ false discovery rate and sized relative to the normalized enrichment score. Only positive associations are shown.

**Figure S12:**
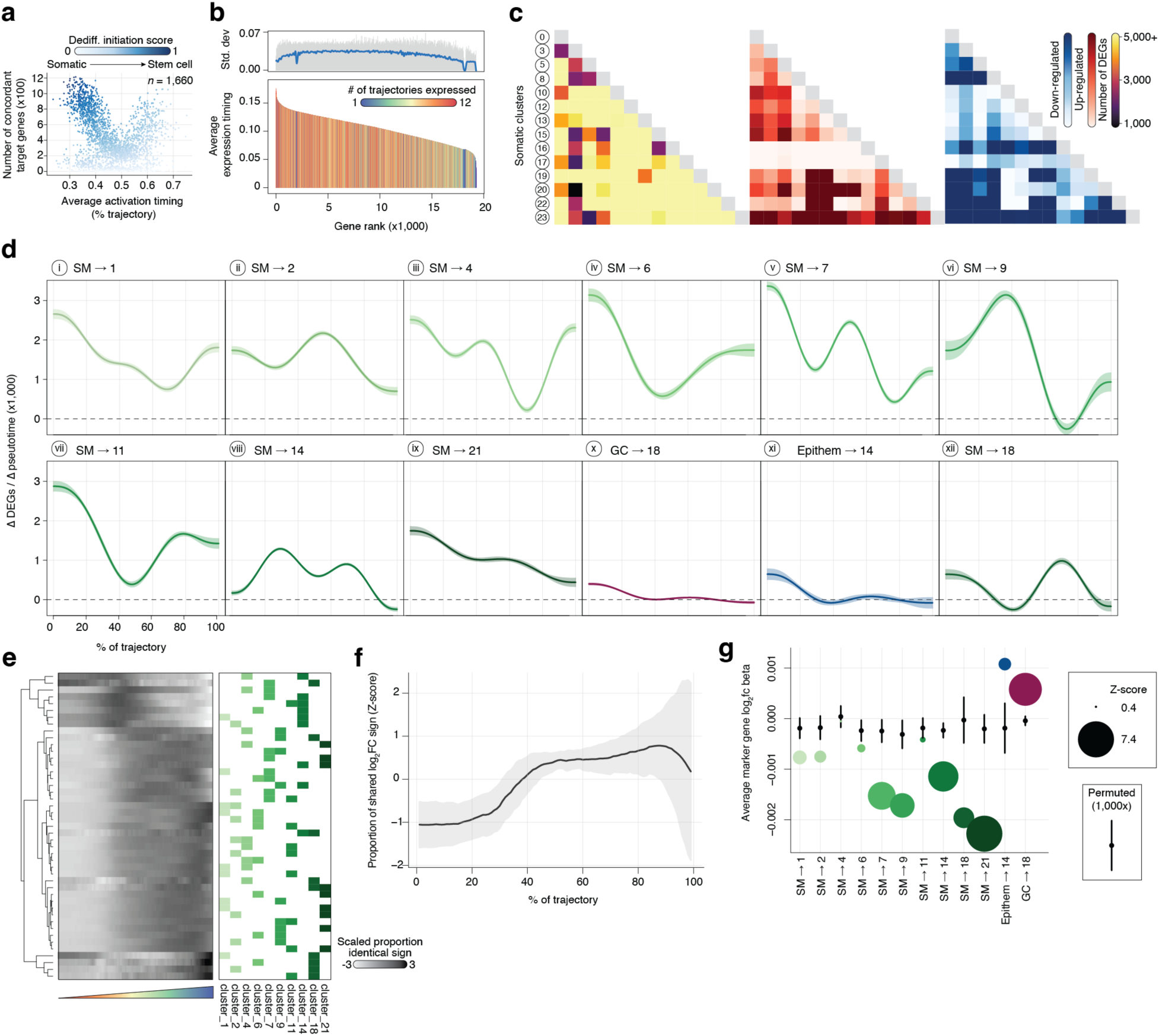
Patterns of conserved and heterogeneous transcriptional programs across diverse somatic dedifferentiation lineages. (**a**) Scatter plot of the number of concordant target genes across trajectories for putative regulators (y-axis) and regulator activation timing (x-axis). Regulators are colored by their dedifferentiation initiation score. (**b**) Ranking of genes by average expression timing across trajectories. Genes are colored by the number of trajectories in which they are expressed. Top, exact (grey bars) and smooth (blue line) standard deviation of expression timing per gene. (**c**) Heatmaps illustrating the number of all (left), up-regulated (middle), and down-regulated (right) differentially expressed genes (DEGs) between all pairwise clusters composed predominantly of cells with somatic identity. (**d**) Rate of change (first derivative) in DEGs vs. T0 at each binned timepoint across dedifferentiation trajectories. Shaded colors indicate the 95% confidence interval. (**e**) Heatmap of row standardized proportion of genes with identical log_2_ fold-change signs at identical pseudotime points for all pairwise trajectories. (**f**) Averaged standardized proportion of genes with identical log_2_ fold-change signs at individual pseudotime points across all trajectories. (**g**) Comparison of mesophyll (green), epithem (blue), and guard cell (purple) marker gene log_2_ fold-changes betas (trend across the trajectory) with randomly permuted (x1,000) non-marker genes.

